# High Precision Binary Trait Association on Phylogenetic Trees

**DOI:** 10.64898/2025.12.24.696407

**Authors:** Ishaq O Balogun, Christopher P Mancuso, Tami D Lieberman

## Abstract

Traditional methods for identifying associations between genomic features and traits, or between pairs of genomic traits, struggle when applied to bacterial genomes. While several microbial GWAS (mGWAS) methods have been developed to account for the fact that genome-wide linkage in bacteria creates strong evolutionary-induced associations, these methods have high false discovery rates or lack statistical power, have poor performance on negative interactions, and face computational limits at the scale required for pangenome-wide study of gene-gene interactions. Here, we present SimPhyNI, a computationally optimized framework for efficient and rigorous mGWAS studies. SimPhyNI builds null co-occurrence distributions by independently simulating traits using phylogenetically-informed parameters, novelly including time to first event. The constrained variation in these simulations, combined with log odds ratio scoring for comparing across traits, robustly identifies both positive and negative associations. Using synthetic datasets mimicking both gene-gene and gene-trait associations, we demonstrate that SimPhyNI achieves high precision and recall for both positive and negative interactions. We demonstrate SimPhyNI’s utility by detecting interactions between phage defense systems in *E. coli* and gene-gene interactions across the entire *E. coli* pangenome (>9 million tests). Though developed here for binary traits, SimPhyNI’s design supports extension to multi-state and continuous traits using generalized models of stochastic simulation. SimPhyNI’s performance and scalability enable genome-wide discovery of genetic interactions that drive microbial function, ecology, and disease.

**Impact Statement:** Understanding how bacterial genes associate with traits and with one another is essential for predicting disease outcomes, antibiotic resistance, and future evolution. However, identifying these interactions is challenging because shared ancestry creates false correlations. SimPhyNI overcomes this through an ancestry-informed statistical simulation process, achieving near-zero false positive rates while maintaining computational efficiency for large scale analyses. This efficiency enables systematic mapping of gene-gene interaction networks across large datasets containing thousands of genes and genomes. As microbial genomic datasets continue to expand, SimPhyNI’s scalability and precision will accelerate discovery of the mechanistic principles underlying infectious disease, microbiome function, and microbial evolution and ecology.

## Introduction

Identifying associations is a critical step in genomic analysis—both for understanding how genetic variation affects traits and how interactions between genomic loci (epistasis) shape evolution. In particular, each bacterial genome contains only a fraction of the genes in its species pangenome [1–3], and understanding the consequences of this variation for health and the gene-gene interactions that constrain evolutionary trajectories are foundational challenges in microbiology [4, 5].

However, traditional genome-wide association studies (GWAS), a standard approach for identifying genetic associations in eukaryotes [6], are not easily applied to bacterial genomes [7]. Microbes reproduce asexually, with limited recombination to break up genes on distant parts of the genome; as a consequence, variants across the genome can be inherited together and perfectly correlate with one another as an artifact of evolutionary relatedness. This non-independence breaks the assumptions of traditional statistical tests, and makes it difficult to define genomic variants and traits that are associated due to true biological interactions [7, 8].

Identifying meaningful biological interactions in microbial genomes requires identifying cases where trait-variant or variant-variant pairs are frequently found together as a result of multiple independent evolutionary events. Traditional statistical methods that ignore phylogenetic relationships cannot differentiate between scenarios with identical co-occurrence counts but different evolutionary histories. For example, a single horizontal gene transfer (HGT) event followed by vertical inheritance may produce the same numerical association as multiple independent events across distant lineages, but the latter pattern provides stronger evidence of mechanistic synergy driving gene co-occurence (Figure 1A). However, reconstructing these evolutionary histories is challenging. Ancient events may be obscured by subsequent losses, horizontal gene transfers, and phylogenetic uncertainty. Methods must therefore balance the need to account for shared ancestry while remaining robust to incomplete evolutionary information.

**Figure 1:**
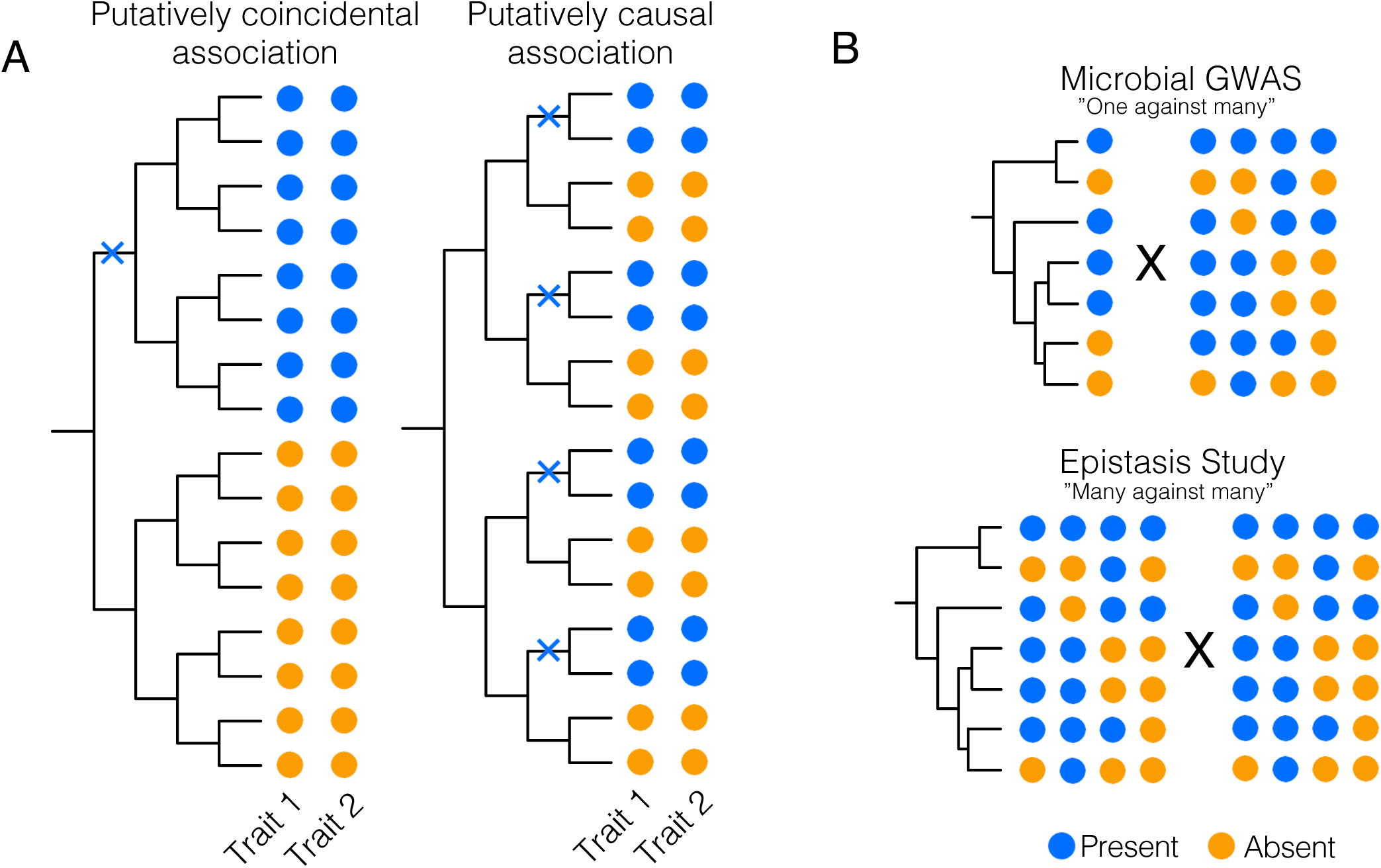
Microbial genome-wide association studies (mGWAS) and epistasis tools aim to correct for phylogenetic signal. (A) Two evolutionary scenarios showcasing the same number of trait cooccurrences, but with different assortment on the same phylogenetic tree. Inferred locations of co-incident trait events are labeled with Xs. While both scenarios are numerically identical for traditional statistical methods that do not consider evolutionary relationships, convergent patterns of co-occurrence across district evolutionary events provides more evidence of a direct relationship between two traits. (B) Microbial GWAS (top) uses a one against many approach to test a single phenotype against an array of genotypic information, yielding a number of comparisons linear with the number of genotypes. In contrast, epistasis studies (bottom), compare sets of genotypic information in a many against many fashion, resulting in a number of comparisons that grows quadratically with the number of genotypes.

Although efforts have been made to incorporate phylogenetic information into association studies [8–10], current methods for microbial GWAS (mGWAS) lack statistical power in common scenarios or struggle to scale to large numbers of statistical tests [7, 11]. Pagel’s correlation method [9] is well established as a standard for the association testing of binary traits; however, it requires fitting computationally expensive statistical models for the likelihood of independent vs dependent evolution. Modern mGWAS tools such as FastLMM [10], TreeWAS [8], and Scoary [12, 13] employ more computationally efficient and widely applicable options, but do not have strong separation between false positives and true associations, creating a tradeoff between stringent false positive control and the recall of interactions. Epistasis studies are particularly challenging for mGWAS studies (Figure 1B), requiring additional statistical power and computational efficiency to evaluate the sheer number of possible pairwise interactions.

Here, we present SimPhyNI (<u>Sim</u>ulation-based <u>Phy</u>logenetic i<u>N</u>teraction Inference), a computationally efficient and robust tool to evaluate associations of binary traits across microbial genomes. SimPhyNI uses the phylogenetic tree to parameterize each trait’s evolutionary history—including rates of gain, loss, and emergence timing—then simulates their independent evolution to estimate expected co-occurrence patterns under a null model of no interaction (Figure 2). Rather than allele frequencies, we focus on binary traits (gene presence/absence) which provide sufficient statistical power with hundreds to thousands of genomes—the current practical limit for rigorous phylogenetic reconstruction [14].

**Figure 2:**
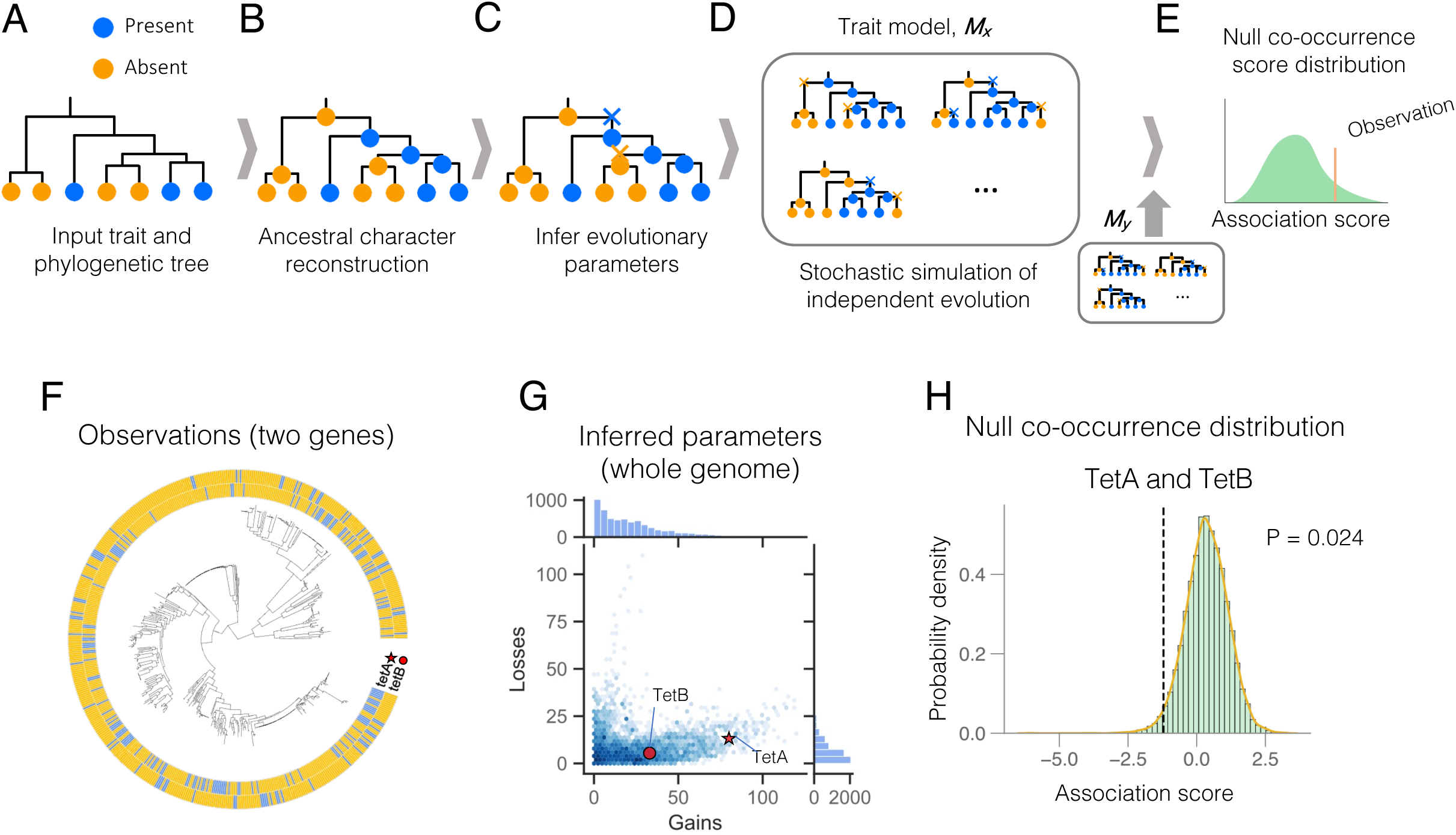
SimPhyNI is a trait association method for mGWAS and Epistasis studies. (A) SimPhyNI begins with an input phylogeny and annotations of trait presence (blue) or absence (orange) on the tips of the tree. (B) Ancestral character reconstruction of the input tree is then performed, using a maximum likelihood approach to infer presence or absence at each node (Methods). (C) Evolutionary parameters of trait gain and loss are inferred using a Markov chain learning process (gain rate, loss rate, and time to first event; Methods). Orange and blue Xs represent inferred state transitions. (D) SimPhyNI then performs many stochastic simulations of a trait under a model of independent evolution using the inferred parameters, resulting in a trait matrix *M_x_* (Methods). (E) This is compared with a trait matrix for the second trait, *M_y,_*, to generate a null co-occurrence score distribution which is compared to the co-occurrence score observed (Methods). (F-H) An example of two interacting genes. (F) Observed presence and absence of tetracycline efflux pump genes, TetA and TetB, on an 500 isolate *E. coli* phylogenetic tree. (G) Distribution of inferred gains and losses for 4333 *E. coli* accessory genes, with TetA and TetB labeled. (H) Histogram of the final statistical readout of TetA and TetB, comparing the observed co-occurrence score with the simulated null distribution. The orange line indicates the approximated distribution (Methods) used for p-value computation.

SimPhyNI achieves superior performance through two key innovations: simulations that account for uncertainty in deep ancestral branches, improving our null models for trait evolution, and a prevalence-aware scoring function, improving statistical power across variable tree topologies. For computational efficiency, we employ kernel density estimation to approximate null distributions, reducing the number of simulations needed to evaluate traits while maintaining statistical power. Together, these optimizations enable SimPhyNI to scale to millions of pairwise interactions without sacrificing statistical rigor or recall.

Using synthetic datasets, we demonstrate that SimPhyNI recovers both positive and negative associations with high accuracy while maintaining exceptionally low false positive rates. We apply SimPhyNI in both a highly directed search for interactions between phage defense systems in *E. coli* and a comprehensive pangenome analysis spanning 9.4 million pairwise associations. As microbial genomic datasets continue to expand, SimPhyNI’s scalability and accuracy will accelerate discovery of the evolutionary constraints and functional dependencies that shape bacterial adaptation and pathogenesis.

## Methods

### SimPhyNI Implemention

#### Trait Parameter Estimation

The first step of SimPhyNI is to estimate the parameters of past trait evolution, including the gain rate, loss rate, the ancestral state, and time to first even for each trait. These values are later used for null simulations. SimPhyNI estimates these parameters using ancestral character reconstruction (ACR), starting with a rooted phylogenetic tree and tip annotations for all traits to be studied (Figure 2A). ACR is performed for each trait independently (Figure 2B) using PastML (1.9.40) with the JOINT maximum likelihood reconstruction method and the F81 character evolution model [15].

To turn ancestral states to parameters that describe the rates of gain and loss for each trait, each phylogenetic tree is traversed from root to tip, recording the minimum number of state transitions between parent and child nodes (Figure 2C). First, the traits state at the root of the phylogeny (root state) and evolutionary distance from the root to the first inferred emergence of each trait (*d*_emergence_(*x*), e.g. first gain or first loss) are calculated. Transition rates are computed by normalizing observed state changes against the total available branch length, expressed in arbitrary evolutionary distance units. Specifically, the gain rate was defined as:

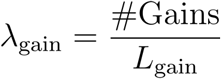

Where *G* is the number of observed trait gains and *L*_gain_ is the total branch length (evolutionary time) during which the trait was absent and available to be gained. Similarly, the loss rate was defined as:

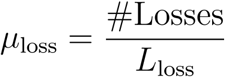

Where *L* is the number of observed losses and *L*_loss_ is the total branch length during which the trait was present and available to be lost. To avoid artificially deflating transition rates due to long branches with hidden transitions, we detected long branches using an interquartile range (IQR) method, excluding those longer than the *Q*_3_ + 0.5 · (*Q*_3_ − *Q*_1_).

#### Trait Model Generation

SimPhyNI then simulates the independent evolution of each trait along the phylogeny as a stochastic process, leveraging the learned parameters from the previous step (Figure 2D). For each trait *x*, this procedure generates a trait model: a set of simulated trait occurrences, represented as a matrix

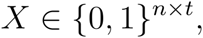

where *n* is the number of terminal nodes (isolates) and *t* is the number of independent simulations.

Simulations are generated using a Markov process that traverses the phylogeny from root to tips. The root state is initialized according to the ACR inferred state. For each branch *b* of length *l_b_*, state transitions are modeled as a Poisson process with transitions are sampled according to gain rate, *λ*, and loss rate, *μ*. If the parent state is absent, the probability of a gain is

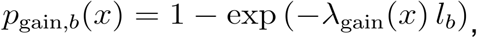

and if the parent state is present, the probability of a loss is

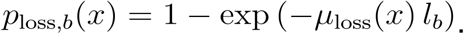

In order to prevent trait emergence at much deeper branches of the tree than empirically observed, evolutionary events are not permitted before nodes that have a cumulative distance from the root less than the emergence threshold *d*_emergence_(*x*) (See Table S1 for overviews of tested variations of this implementation). The results are then stored as a single column matrix *X.* This process is repeated 50 ≤ *t* ≤ 1000 times, depending on whether a coarse or highly resolved approximation of downstream probability distributions is required (see Figure S2-3 for number of comparisons required).

#### Computing Pairwise Co-occurrence Null Distributions

A null distribution of trait co-occurence between pairs of traits is then computed by comparing two trait model matrices (Figure 2E). Let *x* and *y* be two traits of interest, with simulated trait matrices

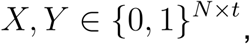

where *N* is the number of terminal nodes and *t* is the number of independent simulations of the trait.

We then consider sets of columns between these two matrices. For each pair of simulation columns (*X_i_*, *Y_j_*), we compute a co-occurrence score using the log odds ratio of tip annotations:

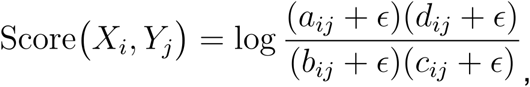

Where:

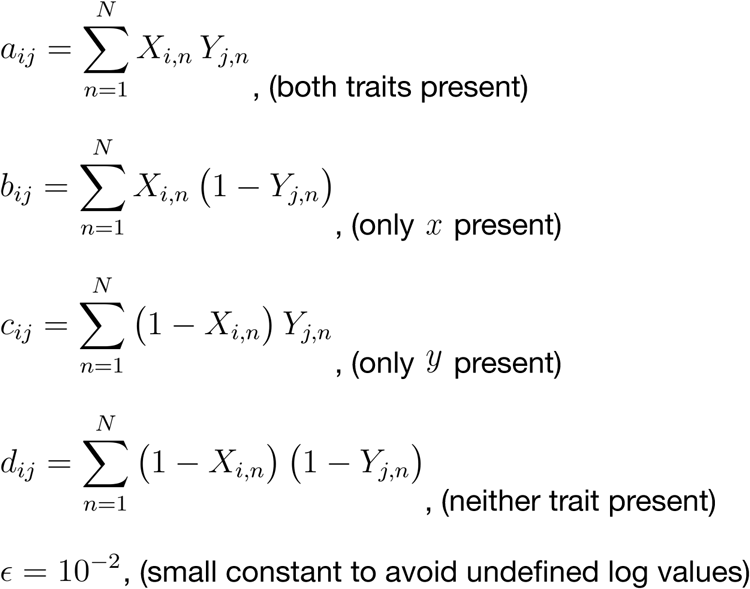

This log odds scoring scoring function was chosen after comparison to other scoring metrics (see Figure S1 and Table S2).

Exhaustively computing scores across all pairs of columns creates a discrete null set of size *t*^2^.

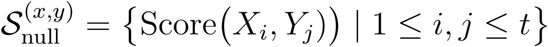

Gaussian kernel density estimation (KDE) is then used to approximate a continuous null distribution of co-occurrence scores from the discrete distribution.

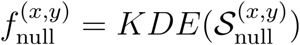

Using estimated null distributions reduces the number of trait simulations required, increasing computational efficiency while preserving statistical power and enabling higher resolution p-values. Silverman’s selection method is applied to determine optimal KDE bandwidths, as observed null distributions are approximately normal (Figure S4; though this approach also handles multimodality sometimes seen, Figure S5). Notably, taking the logarithm of the odds ratio (rather than using the raw odds ratios) is critical for KDE performance, as gaussian KDE requires distributions without high-skew.

#### Calculating Significance and Effect Size

Using the approximated continuous null distribution for each set of traits, 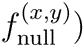, we perform statistical hypothesis testing to identify significant trait-trait associations. For each observed co-occurrence score, an empirical p-value is computed using the approximated distribution (Figure 2E; full example traits Figure 2F-H).

To control for false positives in our biological analyses, we use the Benjamini-Hochberg [16] correction as our default, as it effectively controls the false discovery rate while maximizing recall of interactions. For large-scale pangenome analyses, however, we use the more conservative Benjamini-Yekutieli correction [17]. This is appropriate in this context because it accounts for arbitrary dependencies between tests, such as the co-occurrence of many genes within the same mobile element, and helps filter through the resulting large network of interactions.

Effect sizes were computed as the normalized difference between observed and expected co-occurrence scores:

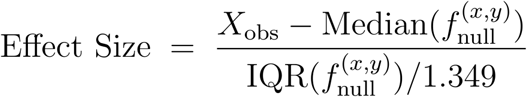

This calculation used median and IQR with a scaling factor in order to resemble Z-score for normal null distributions. However, moderate skew and multimodality have been observed in empirical null distributions (Figure S4,S5), so median and IQR were selected to reduce the sensitivity of effect size to outlying data.

### Synthetic Dataset Construction

Synthetic datasets with varying tree topologies and effect sizes were constructed for the validation and benchmarking of SimPhyNI against existing mGWAS methods. Each dataset consists of a single 500 strain synthetic phylogenetic tree generated using msprime [18] using default parameters and a collection of synthetic interacting trait pairs. For each dataset, we simulated 300 positively correlated pairs, 300 negatively correlated pairs, and 3000 non-correlated pairs of traits; these unbalanced classes emulate the expectation that most gene pairs do not interact.

Synthetic trait pairs were generated using a continuous-time markov process, using the matrix, *Q*, to transition between four states, {00, 01, 10, 11}, representing the union of possibilities of two binary traits (Full Implementation in Supplementary Text 3).

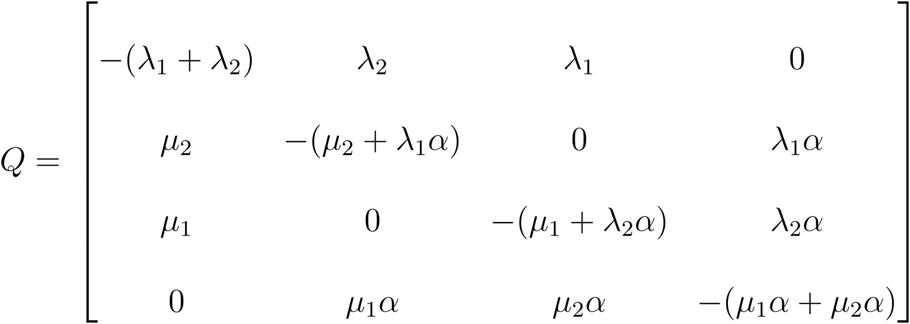

For each trait, gain rates (*λ*), loss rates (*μ*), and root states were jointly sampled from a kernel density estimate of trait parameters from the *E. coli* accessory genome (prevalence 5%-95%) after ACR as described above (Figure S6; see Analysis of the *Escherichia coli* Accessory Genome for dataset used). In order to apply sampled gain and loss rates to new tree topologies—with different total branch sizes, time scales, and numbers of strains—rates were scaled by the ratio of the average non-outlier (see Trait Parameter Estimation) branch length of the tree to be simulated divided by the average non-outlier branch length of the *E. coli* tree used in our pangenome analysis (from which rates distributions were derived).

The interaction parameter is defined as *α* = 10*^I^*, *I* ∈ [-3, 3]. For synthetic data generation *I* takes discrete and decimal values with *I* < 0, *I* > 0, and *I* = 0 yielding negative associations, positive associations and independent evolution, respectively.

Using 11 datasets (each one tree with 3600 synthetic trait pairs) at tested interaction strengths *I* = [− 3, − 2, − 1, −. 75, −. 5, −. 25, 0,. 25,. 5,. 75, 1, 2, 3], the distribution of measured effect sizes was compared to the effect size distribution from SimPhyNI analysis of the *E. coli* pangenome. *I* =− 0. 75 for negative associations and *I* = 0. 75 for positive associations best mirrored effect sizes from the empirical data (Figure S7) and were selected as the standard interaction parameters for method evaluation.

### Benchmarking Against Other Methods

SimPhyNI was benchmarked against several existing methods, including TreeWAS (1.1) [8], FaSTLMM [10] implemented in pyseer (1.3.0) [3], Pagel’s correlation method as implemented in the fitPagel function in the phytools R package (2.4-4) [9], Scoary2 (0.0.25) [12], and Fisher’s exact test, a phylogenetically unaware statistical baseline for binary trait association (Figure 3). All tools were run using the default or recommended parameters specified in their respective publications.

**Figure 3:**
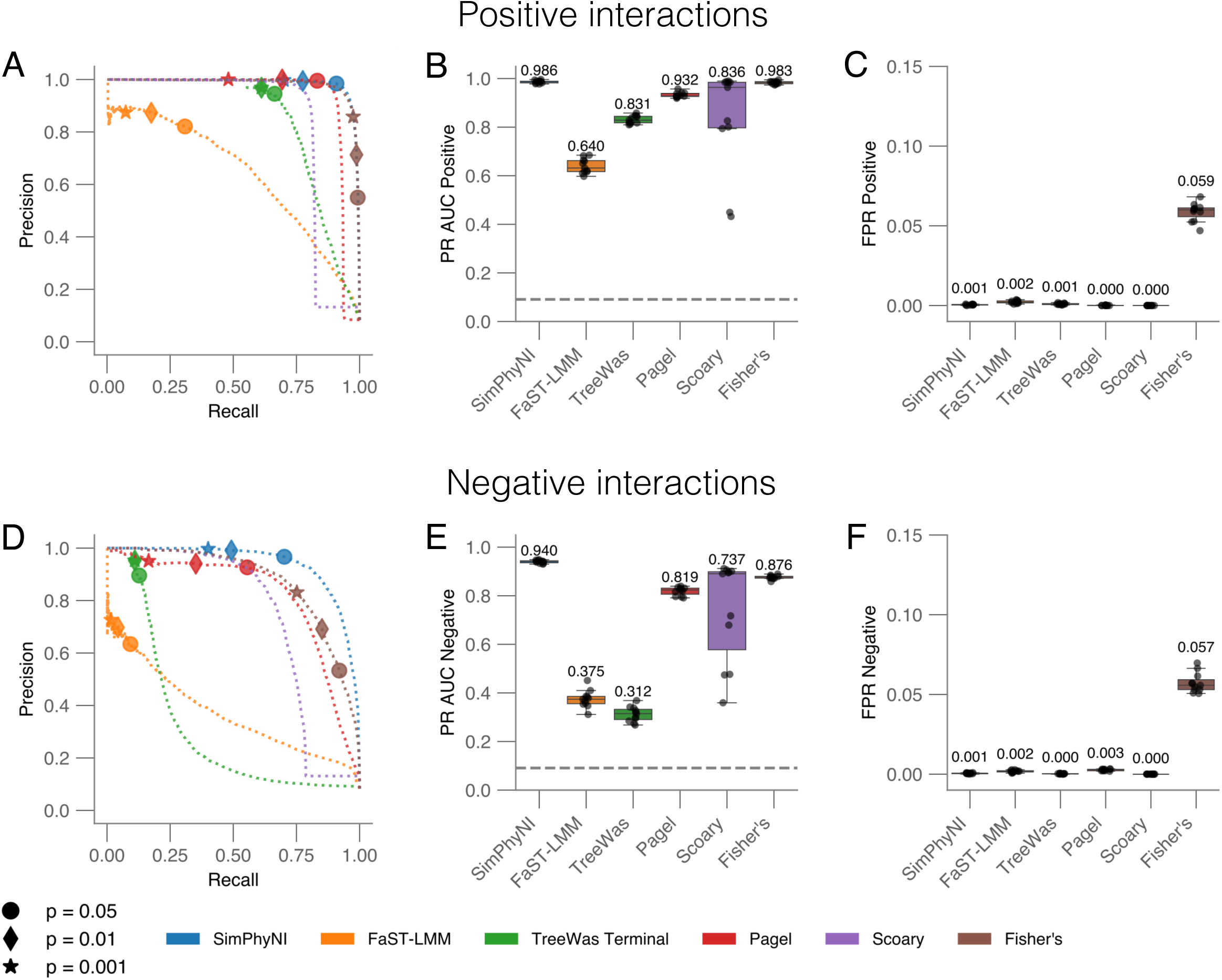
SimPhyNI shows superior performance when compared to other methods using synthetic data. We compared SimPhyNI to 5 other approaches (Methods) for both positive (top row) and negative (bottom row) associations. Each method is in a different color, indicated at the bottom of the figure. (A,D) Precision-Recall curves for tested methods. False Discovery Rate (FDR) adjusted p-value thresholds of 0.05 (circles), .01 (diamonds). and 0.001 (stars) are indicated for each method, except for Scoary which uses a non-p-value based ranking metric. Curves are calculated from the union of 11 synthetic datasets (11 synthetic trees, 3,600 interactions each) with a 1:1:10 ratio of positive:negative:no association pairs. (B,E) Precision-Recall area under curve (AUC) values for each method using 11 synthetic datasets. Box plots show interquartile range and full range across tree typologies. Values closer to 1 indicate better performance, while values closer 0.09 indicate near random performance (dotted line). (C,F) False positive rates (FPR) for each method using 11 synthetic datasets. Box plots show interquartile range and full range across tree typologies. Values closer to 0 indicate better performance. See Figure S8 for performance at different simulated interaction strengths (Methods). See Figure S9 for performance on a differently simulated synthetic data.

Performance metrics for each tool were calculated based on the average of 11 independent synthetic datasets at the tested interaction strength (a total of 11 trees and 39,600 interacting trait pairs; see section Synthetic Dataset Construction). Performance was evaluated at a standardized false discovery rate (FDR) p-value threshold of 0.01 as assessed using Benjamini-Yekutieli using standard classification metrics for positively and negatively associated pairs, including precision, recall, and false positive rate. In addition to threshold-based evaluations, the threshold-independent metric precision-recall area under the curve (PR-AUC) was calculated across reported p-value thresholds (for most methods) or rankings (Scoary).

### Comparative analysis with previously identified *Escherichia coli* phage defense system associations

We re-analyzed a dataset comprising 26,362 *E. coli* genomes previously examined by Wu et al. [19] (Figure 4). This dataset included a neighbor-joining phylogenetic tree and reported annotations of phage defense systems. In their study, Wu et al. employed Pagel’s correlation method with uniform branch lengths (in contrast to the traditional Pagel’s test benchmarked here) to identify significant co-occurrence and exclusion patterns among defense systems, applying a stringent Bonferroni correction (α = 0.05) for multiple testing. The same phylogenetic tree and defense system annotations were input into the SimPhyNI pipeline to compare results across methods. To better align with the statistical conventions for SimPhyNI set by this study, a Benjamini-Hochberg false discovery rate correction was applied at a threshold of 0.05. See Table S4 for full results comparison.

**Figure 4:**
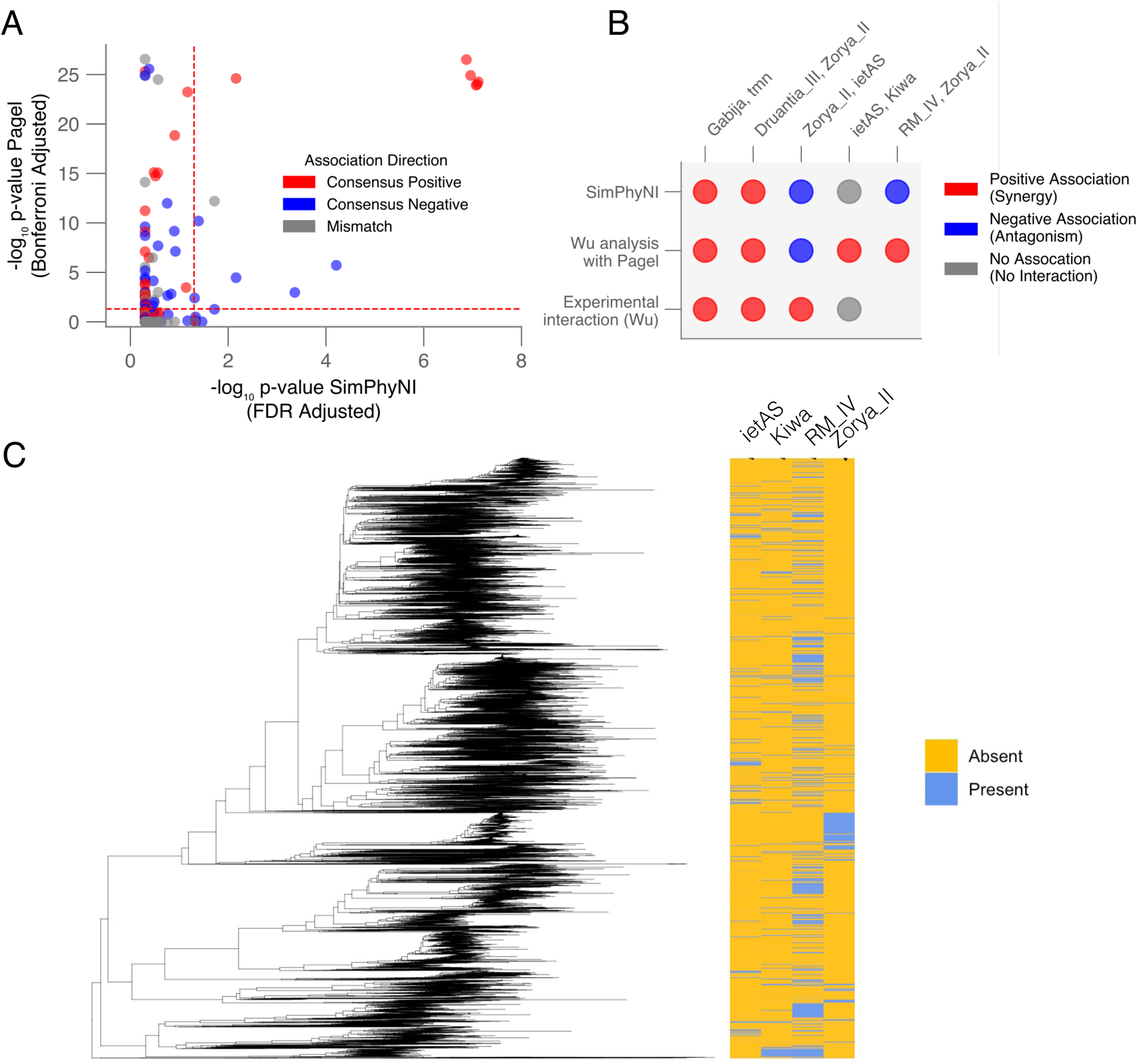
SimPhyNI corroborates known experimentally validated phage defense system associations in *E. Coli*. Wu et al. (2024) annotated phage defense systems, analyzed their co-occurrence using a variant of Pagel’s correlation method, and tested four pairs of systems experimentally. (A) Comparison of p-values from SimPhyNI (FDR adjusted) and Pagel’s correlation method (Bonferroni corrected) for all significant trait associations as determined by either method (Table S4). Red and blue points represent positive and negative associations, respectively, detected by both tools. Gray points indicate system pairs that had conflicting inferred interaction direction between the two methods. Dashed lines indicate significance thresholds after correction. (B) Results of the four experimentally tested interactions ( bottom row, first four columns) compared to predictions from SimPhyNI (top row) and Pagel’s method (middle row). Red indicates a predicted or validated positive interaction, blue indicates a negative interaction, and gray indicates no interaction. One additional gene pair (RM type II and Zorya II) with conflicting directionality despite significance in both tools is shown. (C) The distribution of pairs of systems with conflicted predictions is shown in more detail, with the *E. coli* phylogenetic tree from Wu et al. (2024) on the left and presence and absence of systems indicated on the right.

### Analysis of the *Escherichia coli* Accessory Genome

A representative collection of 500 *E. coli* genomes chosen by PanX [1] and the corresponding maximum likelihood phylogenetic tree were downloaded from NCBI and PanX, respectively. Genomes were annotated using Prokka (1.14.6) [20], and genes were clustered with Panaroo (1.5.2) [2] using the builtin moderate cleaning mode with removal of invalid genes and merging of paralogs. The pangenome output was then filtered for genes between 5% and 95% prevalence. The resulting 4,333 accessory genes were tested for associations using SimPhyNI. The resulting 9.4 million tests were corrected using a 0.01 Benjamini-Yekutieli false discovery rate adjustment (Figure 5, Table S5).

**Figure 5:**
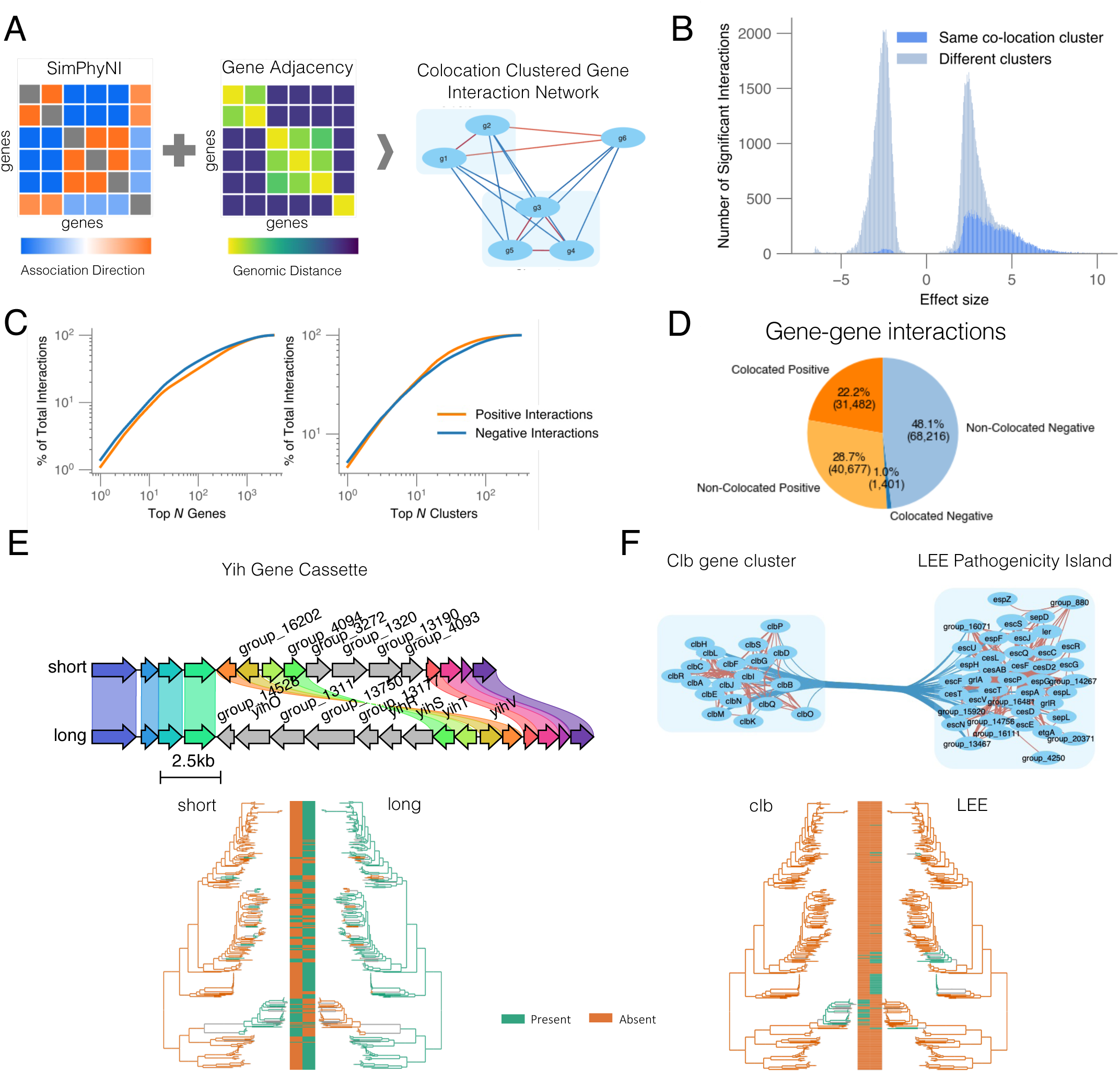
SimPhyNI identifies evolutionary interactions between gene clusters in the *E. coli* pangenome. (A) Gene pairs identified by SimPhyNI (left) that are identified to have shared genomic contexts (middle) are grouped together and assigned to the same co-location cluster (right). (B) Stacked histograms of effect sizes for statistically significant interactions (Benjamini-Yekutieli < 0.01). Gene pairs within the same co-location are indicated in dark blue. Positive effect sizes indicate inferred positive associations; negative effect sizes indicate inferred negative associations; only statistically significant interactions are shown. (C) Cumulative percentage of colocation filtered interactions among top-ranked genes and gene clusters. Genes and clusters are ranked by the number of significant interactions in which they participate (high to low), illustrating that some genes and clusters contribute disproportionately to the interaction network; the 10 top genes and clusters contribute to 10% and 33% of all interactions, respectively. (D) Pie chart indicating the proportion of significant positive and negative gene-gene interactions within and between co-location clusters. (E-F) Two examples of negatively interacting clusters (E) Two structural variants of the yih gene cassette were identified as negatively interacting; an alignment showing homologous flanking regions for both variants, named here ‘long’ and ‘short’ cassettes is shown. Genes unique to each cassette version comprise the clusters and are labeled on the sequence alignment. The phylogenetic tree and heatmap (bottom) show co-occurrence patterns across the *E. coli* population. This negative interaction has been recently described [26]. (F) Interaction network showing negative associations between the clb gene cluster and the LEE pathogenicity island, both associated with *E. coli* virulence, alongside phylogenetic co-occurrence patterns across the *E. coli* population. See Table S5 for all significant interactions.

To remove associations confounded by co-location, a gene graph was built using average genomic distance as edges. Edges were drawn if genes were within 50kb (the scale of a large plasmid). The graph was then clustered using the Leiden graph neighborhood algorithm into 343 gene clusters at a resolution of 2. Varying the resolution from 1 to 2 resulted in only a modest increase in cluster number (326 to 343), indicating that the inferred gene groupings were robust to this parameter choice. Interactions between clusters were aggregated and evaluated for consistency, mean effect size, and density (reported in Table S5). In order to further filter significant interactions, phylogenetic spread for each gene was calculated as the mean of pairwise distances between all genomes in which the gene was present normalized by the maximum possible pairwise distance on the phylogenetic tree (Figure S13).

When evaluating specific cluster–cluster interactions, we assessed negatively associated clusters for conserved genomic context using Clinker (v0.0.31) [21], performing local alignments of complete gene clusters with 10 upstream and 10 downstream flanking genes to identify shared genomic contexts that could explain mutual exclusivity.

## Results

### Development and validation of SimPhyNI using synthetic data

SimPhyNI, like previous microbial trait association methods, is centered around null models of trait-gene and gene-gene co-occurrence derived via simulation [8, 12, 13, 22]. We optimized two key components of this pipeline: 1) how individual traits are simulated, and 2) how trait pair co-occurrences are quantified for each simulation and observation. For trait evolution, we considered Brownian motion and Markov transitions, the two most commonly used models, but selected the latter for its explicit treatment of discrete traits [23].

We found that downstream evaluation metrics were sensitive to Markov model implementation choices (native vs. uniform branch length, including or excluding an emergence threshold, continuous vs discrete time, etc.) (Table S1). Each set of choices produced measurable differences in the distribution of simulated strains across the phylogenetic trees (both prevalence and spread across the phylogeny), resulting in significant effects on detection performance in synthetic data (Figure S1). Similarly, we observed that the choice of scoring function substantially influenced both statistical power and false positive rates (Methods; Table S2, Figure S1).

Therefore, we devised an approach to optimize both of these critical steps. To systematically identify the optimal combination of simulation method and scoring function, we compared performance across sets of approaches using synthetic data. Synthetic datasets were generated using a 4-state continuous-time Markov process [8, 9] with evolutionary parameter rates learned from *E. coli* pangenome genes (Methods). This process was repeated on 11 different simulated phylogenetic trees to ensure the robustness of SimPhyNI across topologies. Pairs of simulated traits were generated for each phylogeny with a class imbalance ratio of 1:1:10 (positive:negative:neutral associations). This class ratio reflects the expectation that neutral associations vastly outnumber true interactions [6, 24]. All method combinations were evaluated using Benjamini-Yekutieli false discovery rate (FDR) correction at a threshold of 0.01.

To perform well in these synthetic datasets a method must (1) maintain high statistical separation of neutral traits associations versus positive or negative associations, (2) display consistent performance across variable tree structures, and (3) recover interactions even after correction for multiple tests. The best-performing configuration combined a log-odds scoring function with a branch length-aware Markov simulation (Methods). This simulation process uses asymmetric gain and loss rates learned from the ancestral character reconstructions (ACR) of traits, then initializes the root of each simulation with the inferred root state ACR and applies a novel parameter, emergence threshold, before which no state changes are simulated (Methods). The combination of simulation method and this scoring function yields exceptional precision-recall area under the curve (PR AUC) values, 0.986 for positive and 0.940 for negative associations, and the low false positive rates (FPR: 0.002 and 0.003) after correction (Figure S1). Given a baseline PR AUC for a random classifier near 0.09 and FPR around 0.9 for this imbalanced classification task, these results reflect strong discriminatory performance. We also note that Markov simulation using normalized branch lengths for tree simulation also performed well across the above metrics, however, normalized simulation had significantly lower recall of both positive and negative interactions.

### SimPhyNI outperforms existing methodologies for trait associations in synthetic data

SimPhyNI was benchmarked against four established association-detection methods developed for microbial traits: Pagel’s correlation [9], FaST-LMM [10], TreeWAS [8], and Scoary [12, 13]. Fisher’s exact test, a basic test for assessing associations between two binary variables, was also included as a baseline method that is not corrected for phylogeny.

Pagel’s correlation is a rigorous likelihood-based framework that compares models of dependent and independent trait evolution on a given phylogenetic tree [9], and only works on binary traits. FaST-LMM [10] and TreeWAS [8] are newer methods designed for broader trait applications, including continuous and multi-state discrete traits. FaST-LMM employs a linear mixed model (LMM) and a kinship matrix to capture the covariance structure among samples [11]. TreeWAS, like SimPhyNI, is simulation-based; however, it differs in implementation details and, more significantly, in the evaluation of trait co-occurrence. TreeWAS natively contains three separate scores for evaluation of interactions; we observed similar trends in evaluation metrics across all three scores and thus only so show the terminal score is shown in Figure 3 (full results in Figure S8). Scoary [12, 13] is a tool for efficient gene presence or absence association testing across a bacterial pangenome. Scoary applies label-switching permutation to filter putative results from Fisher’s exact test. Notably, Scoary uses two distinct metrics for reporting results: a ranking metric (Fisher’s q-value multiplied by the permutation p-value) and a validation metric (permutation p-value alone); we used the ranking metric for PR-AUC calculations and the validation metric for FPR. The same synthetic datasets generated above for optimization were used here for benchmarking across these tools.

Precision-recall analysis demonstrated SimPhyNI’s superior performance compared to these tested methods, achieving average AUC scores of 0.986 and 0.940 for positive and negative associations, respectively, which is significantly higher than all alternative phylogenetic approaches (Mann-Whitney U test p < 10^-4^) except Scoary (Figure 3A,B,D,E). All tested methods consistently outperformed the baseline AUC of 0.09 expected from the class imbalance of the test set. TreeWAS and FaST-LMM had reduced PR AUC values compared to other methods (< 0.85 for positive and < 0.40 for negative interactions) due to their lower recall—likely reflecting their design to prioritize high positive predictive value. Pagel’s correlation showed consistent high separation of neutral and interacting trait pairs across datasets (positive: AUC = 0.932, negative: AUC = 0.819); however, Pagel’s correlation is also the most computationally costly of tested methods by multiple orders of magnitude (Figure S10). Notably the PR AUC of Scoary has substantially higher variance across datasets than all other tested methods, potentially indicating tree topology can affect performance (Figure 3B,E). Fisher’s exact test surprisingly showed high separation in this dataset for positive interactions (AUC=0.983), likely due to the high number of gains and losses for each trait in our simulations (median 29 events per trait); however this discrimination occurred at far below standard p-value cutoffs.

A key strength of SimPhyNI lies in its ability to minimize false positives. Most gene pairs are expected to show no interaction, making stringent FPR control essential [8]. SimPhyNI exhibited consistently near-zero FPRs (<0.001) for both positive and negative associations (Figure 3C,F). All other tested phylogenetic methods have similarly low FPRs. We do note that Scoary—using the permutation test p-value—did not have any tests that were significant after multiple hypothesis correction in any dataset. Though not scalable using standard statistical conventions, we do acknowledge the ability of Scoary to rank hypotheses for potential interaction. Aside from Scoary, the only case that showed significant improvement in FPR over SimPhyNI was Pagel’s correlation for positive associations (p = 0.0008) with all other cases being statistically similar or significantly higher than SimPhyNI. Paired with our analysis of PR AUC, it is clear that SimPhyNI achieved higher statistical separation, retaining low FPRs while recovering more interactions compared to other methods.

All methods detected positive interactions better than negative interactions (Figure 3). This difference likely arises from inherent asymmetries in detecting trait interactions. Negative associations are harder to distinguish from null models than positive associations, especially under low trait prevalence conditions; for example, two traits at 10% prevalence are not expected to overlap much even when non-interacting. Negative interactions between traits lowers their prevalences, further complicating detection. SimPhyNI’s log-odds scoring method helps to overcome this problem by implicitly incorporating prevalence of each trait in its null model (Methods). This adjustment enables robust performance when trait frequencies vary (Figure S1). As a result, SimPhyNI can distinguish genuinely negative interactions from neutral cases nearly as well as it identifies positively associated traits.

We repeated benchmarking across a range of interaction strengths, and found that SimPhyNI consistently outperformed all other tools (Figures S8). We also repeated benchmarking using synthetic datasets generated with directional interactions rather than mutual. We allowed only one trait to influence the transition probabilities of the other, emulating genotype-phenotype associations. Similar performance trends were observed in this dataset, with SimPhyNI outperforming all other tested methods in statistical separation across interaction strengths (Figure S9). This demonstrates that SimPhyNI, though designed for symmetric associations, exhibits strong capability for studying asymmetric relationships such as gene-phenotype or SNP-phenotype interactions.

In addition, SimPhyNI is significantly faster than all other tested methods. We compared computational efficiency using 10 synthetic datasets, each composed of one 500 genome phylogenetic tree, and 1000 traits, analyzing a total of one million trait pairs across tools with standardized compute resources. We observed SimPhyNI to be significantly faster than all other tested methods (Mann-Whitney U test p< 0.01; Figure S10), demonstrating that SimPhyNi’s high precision and recall do not come at the cost of computational efficiency.

### SimPhyNI accurately recovers biologically supported interactions among phage defense systems in *E. coli*

We next sought to demonstrate the utility of SimPhyNI in real-world biological datasets. Prior studies have hypothesized that the evolutionary associations between phage defense systems may be predictive of synergistic defense against phage [19]. To further demonstrate the use of SimPhyNI, we reanalyzed the 26,362-genome *Escherichia coli* dataset from Wu et al. (2024), which computationally inferred synergistic anti-phage activity in *E. coli* and validated a subset of putative interactions between defense systems. The original study employed Pagel’s correlation method with uniform branch lengths (uniform Pagel) to identify co-occurrence and exclusion patterns among phage defense systems. We processed the same neighbor-joining phylogeny and phage defense annotations through the SimPhyNI pipeline.

Across the 2,485 trait pairs tested by both methods, we observed substantial divergence in the significant associations identified (Figure 4A). Only 12 associations were deemed significant by both SimPhyNI and uniform Pagel. Uniform Pagel exclusively identified 50 pairs, while SimPhyNI identified 7 unique associations. Given that Pagel’s method showed highly similar performance to SimPhyNI on the synthetic datasets in this study (Figure 3), the source of this discrepancy remains unclear and may reflect differences in implementation rather than fundamental methodological differences.

Wu et al. experimentally tested four computationally predicted defense system interactions. Two predicted interactions aligned with experimental outcomes: positive associations between Gabija and Tmn as well as Druantia III and Zorya II both showed mechanistic synergy in antiphage activity. However, a predicted negative association between Zorya II and IetAS was also shown to exhibit mechanistic synergy in antiphage activity experiments (Figure 4B). Wu et al. suggested this conflicting case may reflect other evolutionary drivers for the separation of these systems such as metabolic costs or functional redundancy. The fourth prediction, a positive association between IetAS and Kiwa, was found to have no detectable non-additive antiphage activity.

When we applied SimPhyNI to these same interactions, our computational predictions showed stronger concordance with Wu et al.’s experimental findings (Figure 4B). SimPhyNI predicted no interaction between IetAS and Kiwa, which corroborates experimental findings. Further, among the 12 shared significant associations between the two methods, we found one case of directional conflict: the association between RM type IV and Zorya II was called positive by Pagel’s method but negative by SimPhyNI. Visual inspection of the trait distributions on the phylogenetic tree revealed localized patterns of mutual exclusion (Figure 4C), suggesting SimPhyNI’s inference of a negative relationship to be more fitting. These results suggest that SimPhyNI represents a viable and potentially superior alternative approach for large scale trait associations in phylogenetic contexts.

### SimPhyNI enables high-throughput pangenome analysis, identifying biologically meaningful interactions

The power and scalability of SimPhyNI enables analyses that might be computationally prohibitive with existing methods. To demonstrate this capability, we performed a pangenome-wide association screen testing all 9.4 million possible pairs of 4,333 accessory genes across 500 Escherichia coli genomes. This exhaustive pairwise analysis completed in ∼15 minutes when parallelized (64 cores, 64 gb RAM total). SimPhyNI identified 72,159 positive interactions and 69,617 negative associations after multiple-hypothesis correction (Methods).

We hypothesized that large positive effect sizes likely emerged from gene location in syntenic clusters on the genome. To resolve this we constructed a colocation graph from the genomic distances between significant genes pairs and used Leiden clustering to identify communities of co-located genes (Methods, Figure 5A, S11). This resulted in 31,482 positive interactions contained within colocation clusters and, interestingly, 1,401 negative interactions contained within colocation clusters (Figure 5B,D). These negative interactions highlight non homologous genes that have shared genetic neighborhoods, yet rarely exist together in the same genome, likely reflecting mobile elements that compete for integration sites or functional submodules.

When analyzing trends across the *E. coli* pangenome, we observed non-uniform contributions from genes and gene clusters, indicating hubs for evolutionary interactions (Figure 5C). This finding aligns with prior studies that also show hub-like organization of gene-gene associations in the *E. coli* pangenome [25]. However, we noticed some likely false-positive interactions resulting from inferred gains and losses within a poorly resolved ‘outbreak-like’ cluster of the phylogeny resulting in inaccurate trait simulations (Figure S12). We therefore developed a metric for phylogenetic spread (Methods; Figures S13) to sort through interactions alongside effect size. To highlight the increased performance of SimPhyNI on negative associations, we focus on notable examples of predicted antagonism below.

The largest negative interaction between two colocation clusters was also phylogenetically dispersed (mean effect size -4.98). These clusters contain two distinct structural variants of the *yih* gene cassette (Figure 5E). The *yih* gene cassette is responsible for the catabolism of sulfoquinovose, a plant derived sugar used as a sulfur and carbon source, and is present in >95% of *E. coli* within our pangenome study. The structural variants (long and short cassette) differ by a four-gene inversion and unique genetic content; their scattered distribution across the phylogenetic tree suggests frequent recombination (Figure 5E). Their structural and metabolic differences were recently characterized [26], confirming the biological significance of. While previous methods could likely detect the mutual exclusivity of these variants, SimPhyNI’s scalability and prioritization of results through effect size ranking makes this biologically meaningful association immediately evident—something that would be obscured in phylogenetically unaware methods.

SymPhyNI also highlighted interactions that may reflect novel ecological or pathogenic tradeoffs in *E. coli*. We identified a strong negative association between the colibactin (clb) genotoxin island and the LEE pathogenicity island (Figure 5F). These two loci both promote gut colonization of pathogenic *E. coli* and do not share a gene neighborhood (Methods). The clb cluster produces colibactin, a genotoxin that slows epithelial turnover and dampens immune responses [27], whereas the LEE island encodes a type III secretion system that remodels the host epithelium and promotes adherence [28]. Though both confer host-associated advantages, the two islands show within phylogroup mutual exclusivity with an average effect size of -2.37 (Methods). This pattern suggests that maintaining both virulence systems simultaneously is either metabolically costly, functionally redundant, or ecologically disadvantageous—all hypotheses that could be evaluated through future experimental work.

Together, these results illustrate how SimPhyNI can be used to effectively analyze and order large-scale genome-wide data, recover known interactions and provide hypotheses for further mechanistic study within microbes. A list of all putatively interacting gene and cluster pairs in *E. coli* is reported in Table S5 to serve as a resource for the community.

## Discussion

We have shown that SimPhyNI achieves high precision and recall in identifying evolutionary interactions (Figure 3). Notably, it: (1) excels in detecting negative associations (antagonisms), which are frequently missed by other tools; and (2) achieves near-zero false positive rates, which is critical due to the sparsity of true interactions relative to all possible trait pairs. This robustness makes SimPhyNI reliable for exploratory analyses where spurious hits could otherwise overwhelm interpretation. We show multiple use-cases of SimPhyNI’s accuracy, including target searches on large phylogenetic trees (Figure 4) and network recovery with large numbers of genes (Figure 5)

SimPhyNI’s strong statistical performance was enabled by two key innovations: a novel parameter for trait simulation that accounts for uncertainty in deep ancestral branches of a phylogeny and a scoring function that accounts for variation in trait prevalence across simulation (Figure 2).

Moreover, SimPhyNI achieves superior efficiency (Figure S10) using two key innovations. First, stable p-values for co-occurrence comparisons are generated from a small number of simulations (generally 64) using a KDE to approximate null co-occurrence distributions from the combinatorial comparison of simulated traits (Figure S2). Second, though implemented in python, we compile key functions to machine code for space and time efficiency. These improvements enable computational cost to be drastically reduced, thus powering large-scale comparisons that power analysis of genome-wide epistasis (Figure 5).

Although SimPhyNI maintains high precision in binary trait association, detecting interactions in evolutionary data remains inherently challenging. Trait pairs with low prevalence or limited phylogenetic spread may remain undetectable despite true underlying interactions. Further, true of gain and loss events can be invisible to phylogenetic reconstruction—particularly on long branches, reducing the effective information available to infer associations. We have attempted to minimize the impact of these missing events by including a time to first event parameter. In addition, our synthetic data used for testing allows for repeated gain and loss along single branches of the phylogeny to emulate this loss of information (Figure 3,S8,S9). In these simulations, SimPhyNI outperforms other methods, maintaining low false positive rates while detecting more interactions than similar methods—across both diverse tree topologies and simulation models.

We have established that SimPhyNI shows excellent performance for binary traits, which are central to many questions in microbial evolution (including gene presence/absence, binary phenotypes, and disease incidence). Importantly, its architecture is not limited to this setting. The underlying framework is readily extendable to multi-state and continuous traits through larger Markov transition matrices or Brownian motion models, and the current log-odds scoring function could be generalized to multiclass likelihood statistics. These extensions could broaden the scope of SimPhyNI to encompass SNPs, peptide variants, and expression levels, enabling direct analysis of high-dimensional trait data. Thus, while SimPhyNI already surpasses existing methods in binary scenarios, its flexible design positions it as a foundation for future models that accommodates a broader range of microbial trait evolution.

Altogether, SimPhyNI is a robust, extensible framework for evolutionary interaction inference, with computational efficiency that opens new avenues for large-scale exploration of associations and co-evolution in microbial genomes.

## Abbreviations

ACR: Ancestral Character Reconstruction
BY: Benjamini–Yekutieli
BH: Benjamini–Hochberg
FDR: False Discovery Rate
FPR: False Positive Rate
GWAS: Genome-Wide Association Study
HGT: Horizontal Gene Transfer
KDE: Kernel Density Estimation
LMM: Linear Mixed Model
mGWAS: Microbial Genome-Wide Association Study
NJ: Neighbor Joining
PR AUC: Precision–Recall Area Under the Curve
SimPhyNI: Simulation-based Phylogenetic iNteraction Inference

## Contributions

Ideation: I.O.B., C.P.M.,T.D.L.; Tool development: I.O.B.; Computational Validation: I.O.B., Biological Validation: I.O.B., C.P.M.; Original manuscript draft: I.O.B., T.D.L.

## Acknowledgments

We thank all members of the Lieberman Lab for critical advice on this project and feedback on the manuscript and Michael Laub for discussions on phage-defense elements. The authors acknowledge the MIT Office of Research Computing and Data for providing high performance computing resources.

## Data Summary

SimPhyNI is publicly available at https://github.com/jpeyemi/SimPhyNI, and code for related benchmarking, validation, and biological analyses are available at doi.org/10.5061/dryad.9kd51c5xt. The neighbor-joining phylogenetic tree and phage defense system annotations used in this study were obtained from Wu *et al.* (2024). A representative set of *Escherichia coli* genomes and the corresponding maximum-likelihood phylogenetic tree were downloaded from the PanX database (https://pangenome.org/Escherichia_coli).

## References

1. Ding W, Baumdicker F, Neher RA. panX: pan-genome analysis and exploration. Nucleic Acids Res 2018;46:e5.

2. Tonkin-Hill G, MacAlasdair N, Ruis C, Weimann A, Horesh G, et al. Producing polished prokaryotic pangenomes with the Panaroo pipeline. Genome Biol 2020;21:180.

3. Lees JA, Galardini M, Bentley SD, Weiser JN, Corander J. Pyseer: A comprehensive tool for microbial pangenome-wide association studies. Bioinformatics 2018;34:4310–4312.

4. Lang AS, Buchan A, Burrus V. Interactions and evolutionary relationships among bacterial mobile genetic elements. Nat Rev Microbiol 2025;23:423–438.

5. Johnson MS, Reddy G, Desai MM. Epistasis and evolution: recent advances and an outlook for prediction. BMC Biol 2023;21:120.

6. Uffelmann E, Huang QQ, Munung NS, de Vries J, Okada Y, et al. Genome-wide association studies. Nat Rev Methods Primers 2021;1:1–21.

7. San JE, Baichoo S, Kanzi A, Moosa Y, Lessells R, et al. Current affairs of microbial genome-wide association studies: Approaches, bottlenecks and analytical pitfalls. Front Microbiol 2019;10:3119.

8. Collins C, Didelot X. A phylogenetic method to perform genome-wide association studies in microbes that accounts for population structure and recombination. PLoS Comput Biol 2018;14:e1005958.

9. Pagel M. Detecting correlated evolution on phylogenies: a general method for the comparative analysis of discrete characters. Proc Biol Sci 1994;255:37–45.

10. Lippert C, Listgarten J, Liu Y, Kadie CM, Davidson RI, et al. FaST linear mixed models for genome-wide association studies. Nat Methods 2011;8:833–835.

11. Saber MM, Shapiro BJ. Benchmarking bacterial genome-wide association study methods using simulated genomes and phenotypes. Microb Genom;6. Epub ahead of print March 2020. DOI: 10.1099/mgen.0.000337.

12. Roder T, Pimentel G, Fuchsmann P, Stern MT, von Ah U, et al. Scoary2: rapid association of phenotypic multi-omics data with microbial pan-genomes. Genome Biol 2024;25:93.

13. Brynildsrud O, Bohlin J, Scheffer L, Eldholm V. Rapid scoring of genes in microbial pan-genome-wide association studies with Scoary. Genome Biol 2016;17:238.

14. Lees JA, Harris SR, Tonkin-Hill G, Gladstone RA, Lo SW, et al. Fast and flexible bacterial genomic epidemiology with PopPUNK. bioRxiv 2018;360917.

15. Ishikawa SA, Zhukova A, Iwasaki W, Gascuel O. A fast likelihood method to reconstruct and visualize ancestral scenarios. Mol Biol Evol 2019;36:2069–2085.

16. Benjamini Y, Hochberg Y. Controlling the false discovery rate: A practical and powerful approach to multiple testing. J R Stat Soc Series B Stat Methodol 1995;57:289–300.

17. Benjamini Y, Yekutieli D. The control of the false discovery rate in multiple testing under dependency. Annals of Statistics 2001;29:1165–1188.

18. Baumdicker F, Bisschop G, Goldstein D, Gower G, Ragsdale AP, et al. Efficient ancestry and mutation simulation with msprime 1.0. Genetics;220. Epub ahead of print 3 March 2022. DOI: 10.1093/genetics/iyab229.

19. Wu Y, Garushyants SK, van den Hurk A, Aparicio-Maldonado C, Kushwaha SK, et al. Bacterial defense systems exhibit synergistic anti-phage activity. Cell Host Microbe 2024;32:557–572.e6.

20. Seemann T. Prokka: rapid prokaryotic genome annotation. Bioinformatics 2014;30:2068–2069.

21. Gilchrist CLM, Chooi Y-H. Clinker & clustermap.Js: Automatic generation of gene cluster comparison figures. Bioinformatics 2021;37:2473–2475.

22. Godfroid M, Coluzzi C, Lambert A, Glaser P, Rocha EPC, et al. Evo-Scope: Fully automated assessment of correlated evolution on phylogenetic trees. Methods Ecol Evol 2024;15:282–289.

23. Phylogenetic Comparative Methods · learning from trees. https://lukejharmon.github.io/pcm/ (accessed 16 September 2025).

24. Zhou W, Nielsen JB, Fritsche LG, Dey R, Gabrielsen ME, et al. Efficiently controlling for case-control imbalance and sample relatedness in large-scale genetic association studies. Nat Genet 2018;50:1335–1341.

25. Hall RJ, Whelan FJ, Cummins EA, Connor C, McNally A, et al. Gene-gene relationships in an Escherichia coli accessory genome are linked to function and mobility. Microb Genom;7. Epub ahead of print September 2021. DOI: 10.1099/mgen.0.000650.

26. Kaznadzey AD, Rybina AA, Bessonova TA, Korshunov DS, Tutukina MN, et al. Sulfoquinovose Catabolism in E. coli Strains: Compositional and Functional Divergence of yih Gene Cassettes. Int J Mol Sci 2025;26:10351.

27. Auvray F, Perrat A, Arimizu Y, Chagneau CV, Bossuet-Greif N, et al. Insights into the acquisition of the pks island and production of colibactin in the Escherichia coli population. Microb Genom;7. Epub ahead of print May 2021. DOI: 10.1099/mgen.0.000579.

28. Ingle DJ, Tauschek M, Edwards DJ, Hocking DM, Pickard DJ, et al. Evolution of atypical enteropathogenic E. coli by repeated acquisition of LEE pathogenicity island variants. Nat Microbiol 2016;1:15010.

