## Supplemental Figure S1-S13, Table S1-S3 for "High Precision Binary Trait Association on Phylogenetic Trees"

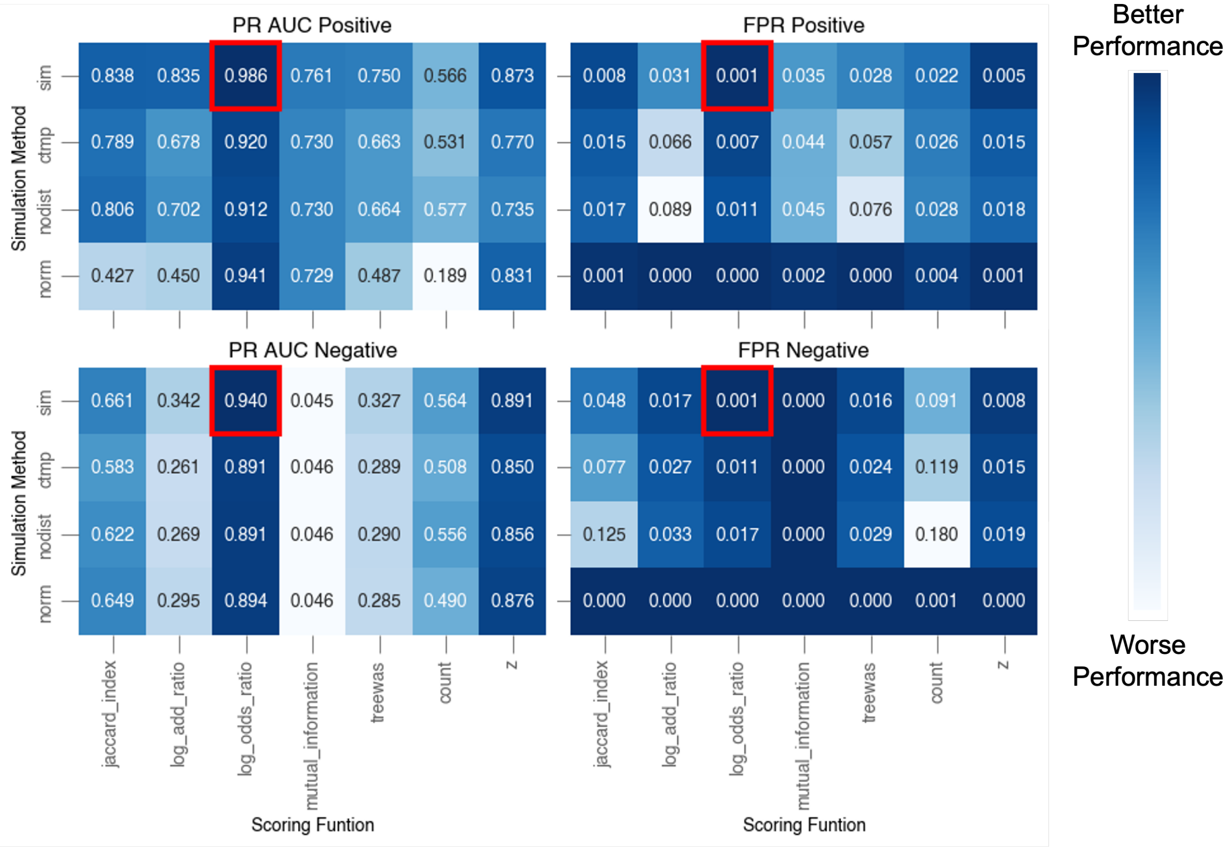


Figure S1: **Metrics for all test statistics and simulation methods using 4-state Markov data generation using Benjamini-Yekutieli correction of 0.01** Each pair of heatmaps reports a performance metric (PR AUC or FPR) across combinations of simulation methods (rows) and scoring function (columns). The top and bottom rows report performance for the identification positive and negative associations, respectively. The log odds ratio scoring function was observed to have the highest performance across metrics. Notably our default simulation method (sim) and normalized simulation (norm) have the best performance, with default simulation having significantly higher positive PR AUC and significantly lower values across other displayed metrics. However the magnitudes of the KS statistics for each metric above paired with the observed higher recall of default simulation under stringent false discovery control highlight this method as a superior branch length normalized simulation.

Table S1: Simulation Methods Implemented in SimPhyNI

| **Simulation Method** | **Implementation Details** |
|---|---|
| `base simulation` | Markov process with a maximum of one event per branch. Uses a distance threshold, before which no events are simulated. |
| `nodist` | `base simulation` implementation without a distance threshold for trait emergence. |
| `ctmp` | Continuous-time implementation of `base simulation` allowing multiple events per branch using exponential waiting times. Also lacks distance threshold, relying on the end state distribution to prevent long ancestral branches from dominating simulation outcomes |
| `norm` | `base simulation` with all branch lengths normalized to a fixed value (1). Uses number of nodes to calculate gain and loss rates |

Table S2: Scoring Functions for Trait Co-occurrence

| **Scoring Function** | **Formula** | **Observations** |
|---|---|---|
| Jaccard Index | $\frac{A \cap B}{A \cup B}$ | Simple similarity metric. Easy to interpret but sensitive to low prevalence. Limited statistical power. |
| Log add ratio | $\log\left(\frac{(A \cap B) + (\neg A \cap \neg B)}{(\neg A \cap B) + (A \cap \neg B)}\right)$ | Emphasizes co-occurrence and co-absence over mismatches. Sensitive to imbalanced marginals. |
| Log odds ratio | $\log\left(\frac{(A \cap B)(\neg A \cap \neg B)}{(A \cap \neg B)(\neg A \cap B)}\right)$ | Statistically powerful. Used in SimPhyNI null model testing. Performs well across prevalence spectra. |
| Mutual Information | $\sum_{A,B \in \{0,1\}} P(A \cap B) \log\left(\frac{P(A \cap B)}{P(A)P(B)}\right)$ | Measures dependency between presence/absence vectors $A$ and $B$. Symmetric. Sensitive to sampling noise. Requires smoothing with small sample sizes. |
| TreeWAS Terminal Statistic | $(A \cap B) + (\neg A \cap \neg B) - (\neg A \cap B) - (A \cap \neg B)$ | Captures agreement between predicted and observed trait values across descendant nodes. Used in TreeWAS to evaluate phylogenetically corrected association strength. |
| Count | $A \cap B$ | Simple co-occurrence count. Does not adjust for background rates. Biased by prevalence. |
| Z score | $\frac{A \cap B - P(A)P(B)N}{P(A)P(B)(1 - P(A)P(B))}$ | Standardized deviation from expected co-occurrence. Can inflate importance with large $N$. |

**Legend:**
$N$ : Total number of genomes or samples.
$A$, $B$ : Binary vectors indicating trait presence/absence across $N$ samples.
$A \cap B$ : Count of samples where both traits are present.
$A \cup B$ : Count of samples where either trait is present.
$\neg A$ : Complement of $A$ (absence of trait $A$).
$P(A)$, $P(B)$ : Proportions of samples with traits $A$, $B$ respectively.
$\epsilon$ (not shown): Small positive pseudocount used for smoothing (typically 1).

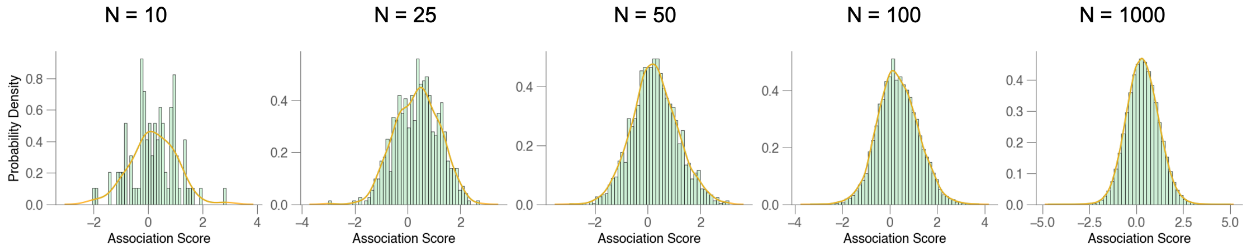


Figure S2: **Comparison of KDE versus discrete null distributions for trait co-occurrence scores.** Presented as as visual for null co-occurrence distribution approximation using kernel density estimation. $N$ represents the number of simulations used for each trait therefore plots show $N^2$ co-occurrence observations corresponding to the exhaustive comparison of trait simulations. As $N$ increases the discrete observations (green bars) and KDE (orange line) converge to the same normal distribution.

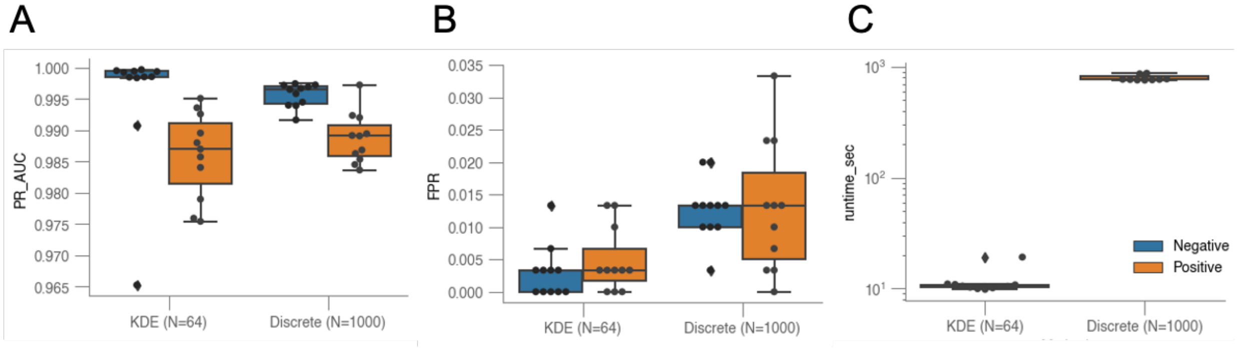


Figure S3: **Comparison of SimPhyNI using KDE and Discrete null models shows improved performance across metrics when using estimated distributions** Comparisons of KDE and discrete null models for SimPhyNI across 11 synthetic datasets generated using 4-state Markov transitions with interaction parameters of $I = [-2, 0, 2]$. Results show statistically significant improvements on PR AUC for negative associations ($p = 0.0125$), FPR for negative associations ($p = 0.000689$), and runtime ($p = 8.15 \times 10^{-5}$) favoring KDE over discrete null models, as well as comparable performance between both methods for PR AUC and FPR in positive associations ($p = 0.138$ and $p = 0.0368$, respectively). P-values were calculated using a two-sided Mann–Whitney U test.

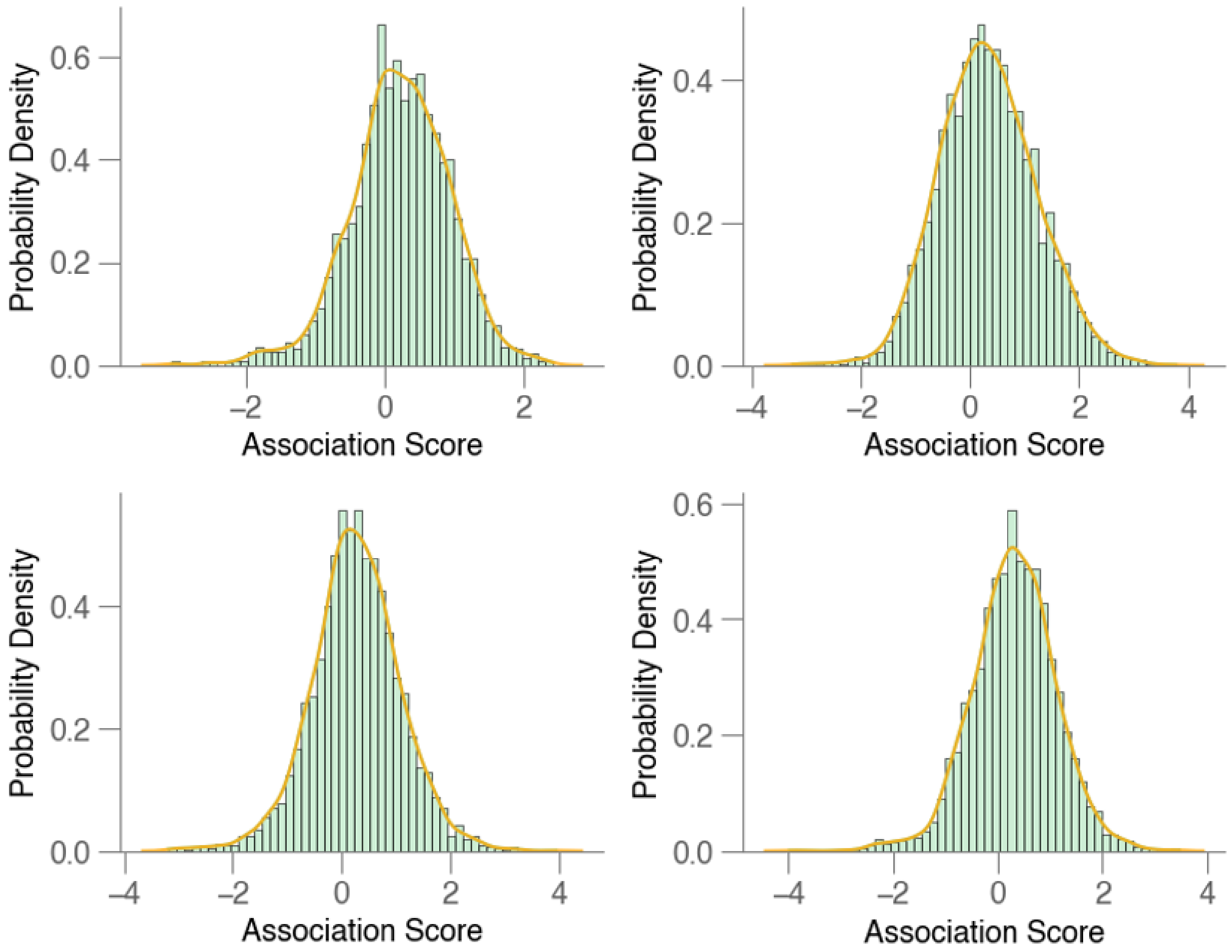


Figure S4: **Example null distributions show visual fit of KDE near normal** Presented as an example of typical null distributions encountered when running SimPhyNI. Distributions are near normal, showcasing minor skew and/or non-zero central tendencies—features well fit by KDE. Distribution approximations are used to determine p-value and effect size for interactions.

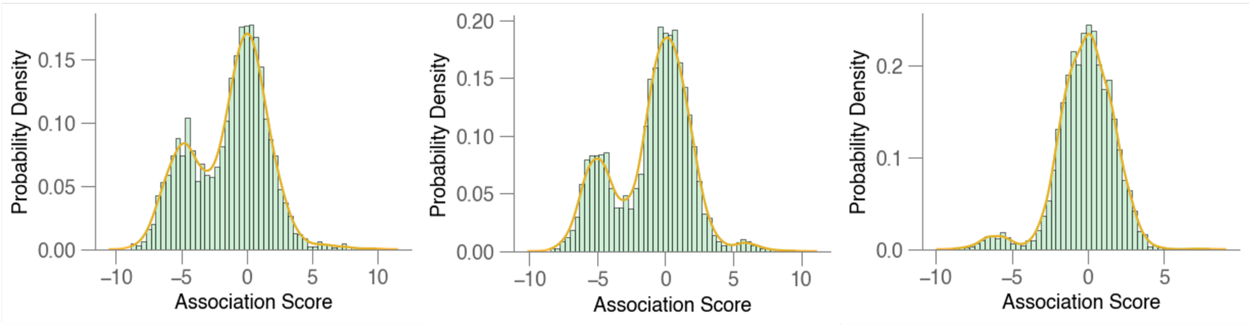


Figure S5: **Example multi-modal null distributions show visual fit of KDE for non-normal data** Presented as an example of multi-modal null co-occurrence distributions that can arise due to a combination of topological features and trait parameters during simulation. KDE with $N = 64$ shows a visually suitable fit to these distributions, capturing non-normal behavior. Each plot is a distinct pair of synthetic traits generated using the 4-State Markov process.

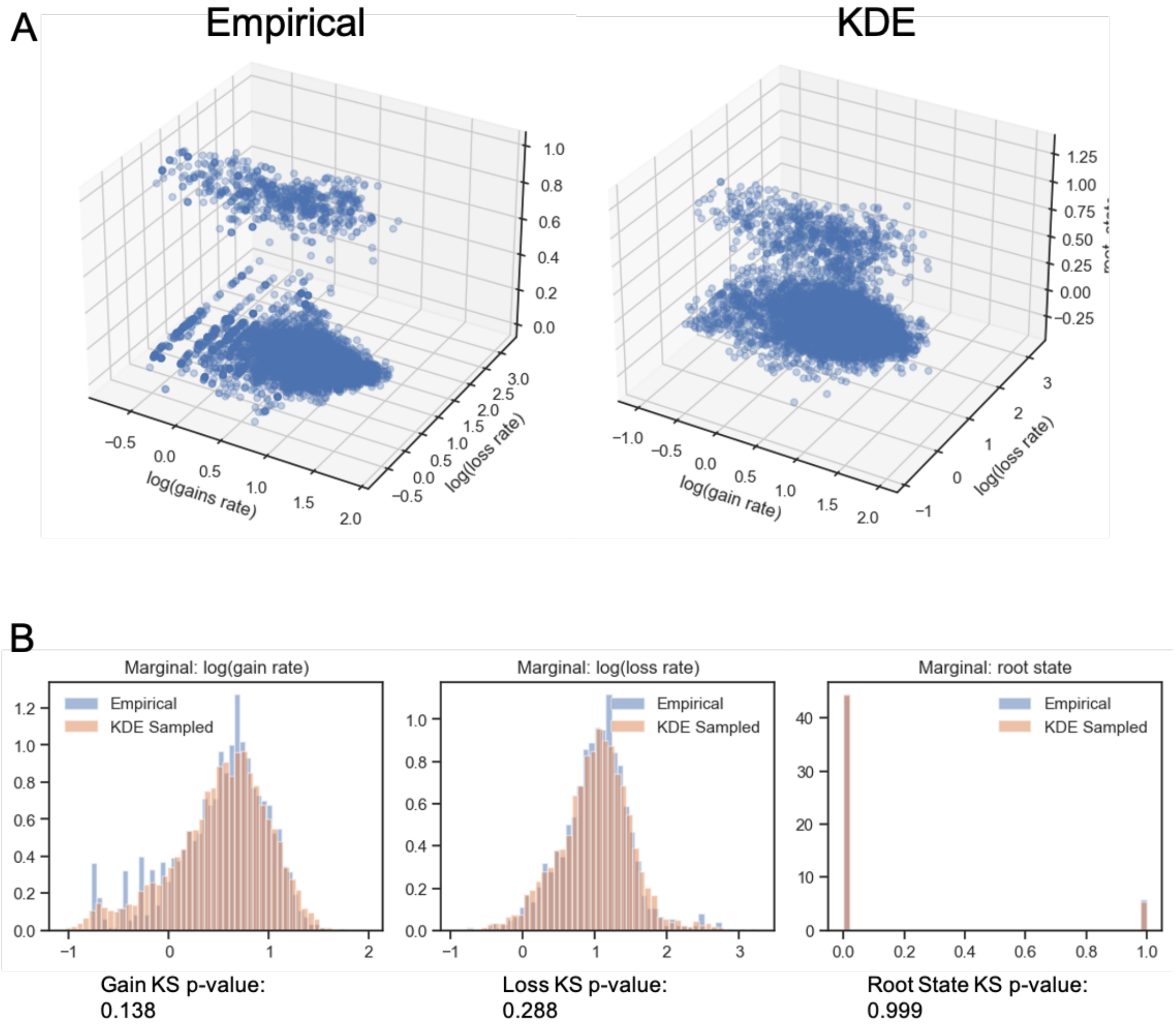


Figure S6: ***E. coli* trait parameter distribution kernel density estimation for sampling in synthetic data.** (A) Empirical gain rates, loss rates, and inferred roots states for *E. coli* 5903 accessory genes and 5903 samples from a 3-dimensional KDE. Root state, of sampled data was rounded to the nearest integer. (B) Marginal plots of each dimension, showing the overlap of empirical and KDE sampled data. Empirical and KDE sampled distributions are non significantly different (Kolmogorov–Smirnov Test, $P > 0.13$), allowing non redundant sampling of trait parameters for synthetic data generation.

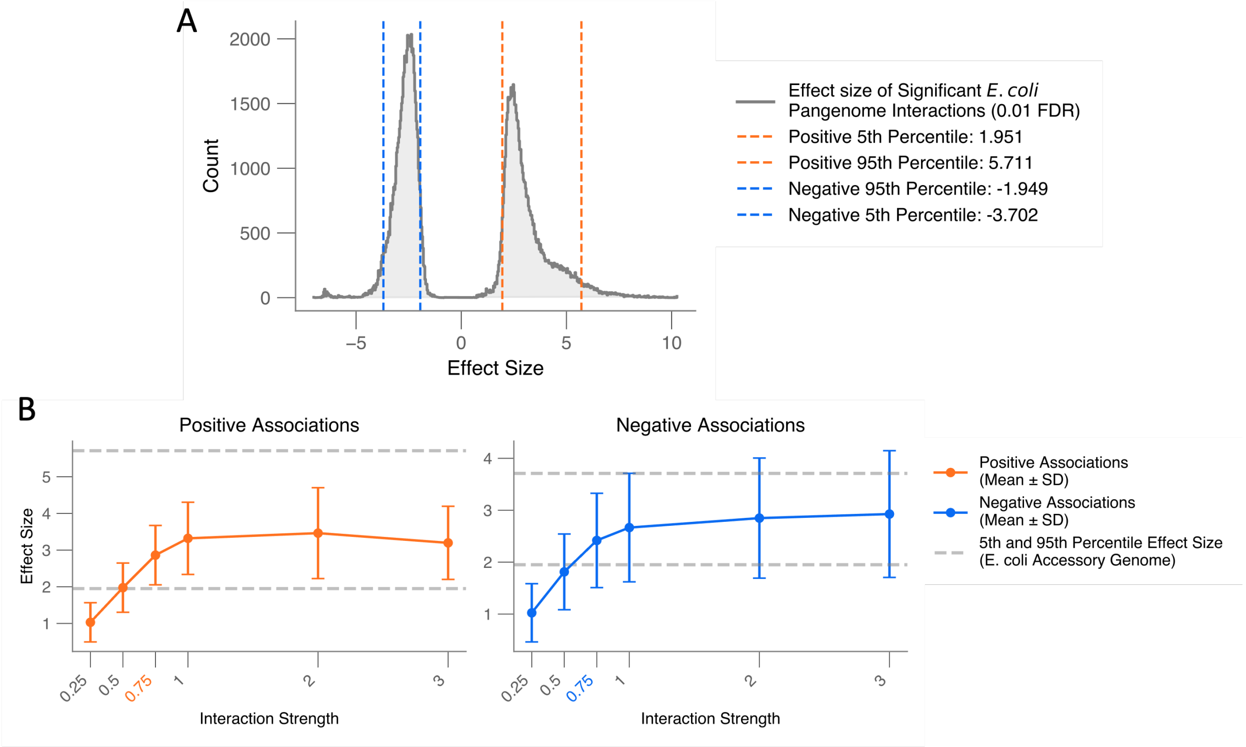


Figure S7: **Distributions and benchmarking of synthetic data input interaction strength against effect sizes from significant trait-trait associations in the *E. coli* pangenome.** (A) Histogram of effect sizes for all significant associations (FDR $< 0.01$) identified by SimPhyNI. Dashed vertical lines indicate the 5th and 95th percentiles of effect sizes for positive (orange) and negative (blue) associations, providing empirical thresholds for interaction effect sizes. (B) Mean $\pm$ standard deviation of recovered effect sizes across varying interaction strengths in synthetic benchmarking datasets. Positive (left, orange) and negative (right, blue) association classes are shown separately. The dashed lines mark the empirical 5th and 95th percentile effect sizes from the real *E. coli* data (A) used as a reference for selecting representative interaction strengths ($I = \pm 1$) for synthetic data evaluation.

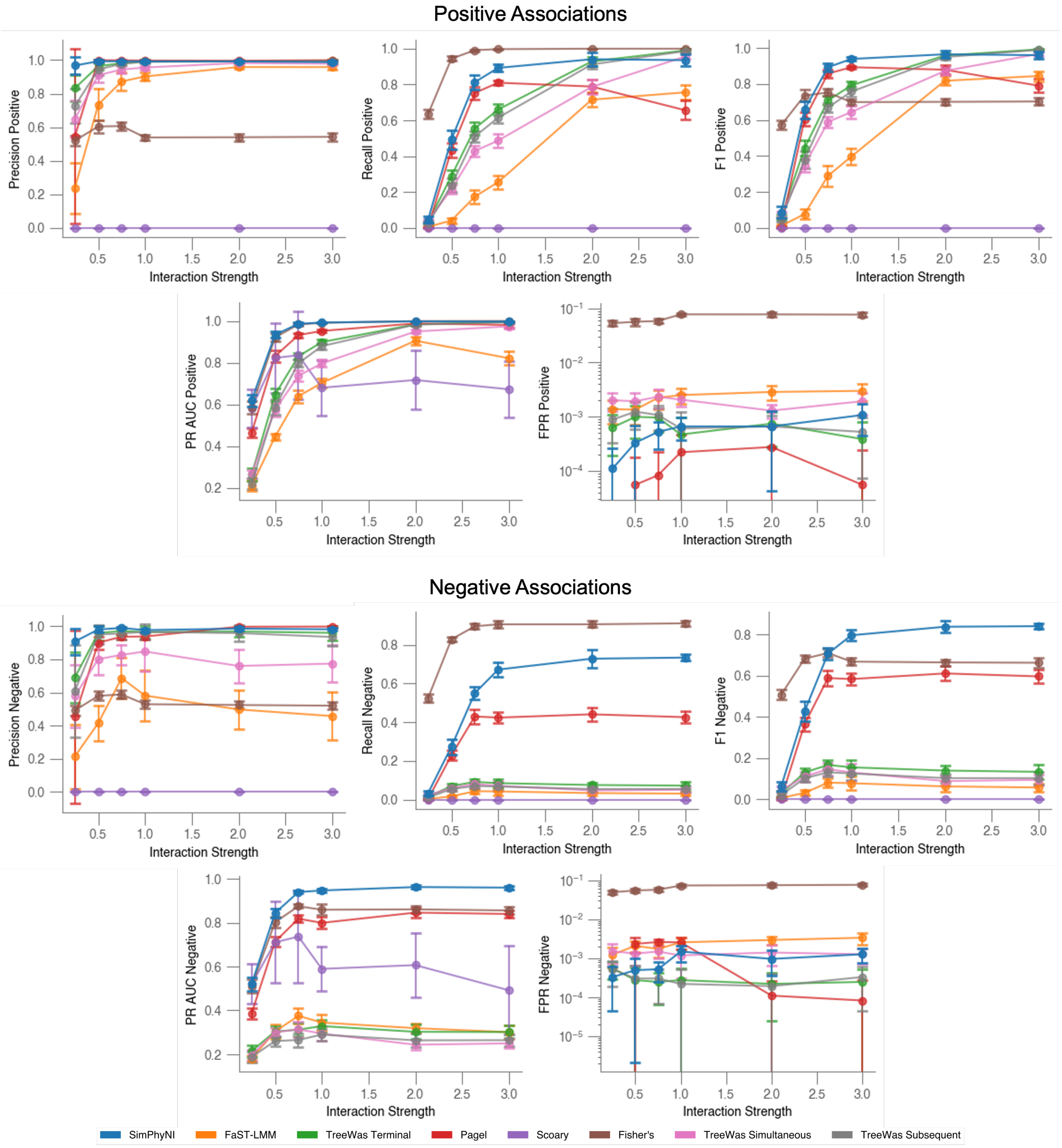


Figure S8: **All metrics for all tested methods using 4-state Markov data generation across all tested interaction strengths.** Benchmarking of all tested methods using synthetic datasets and evaluated using a Benjamini-Yekutieli multiple testing correction at FDR = 0.01. Points show mean ± standard deviation for 11 distinct phylogenetic trees at the given interaction strength. Five metrics are shown: precision, recall, F1-score, PR AUC, and false positive rate (FPR), each for both positive and negative association classes. SimPhyNI outperforms other tools across most metrics, achieving better performance at lower interaction strengths and converging to superior values as interaction strength increases.

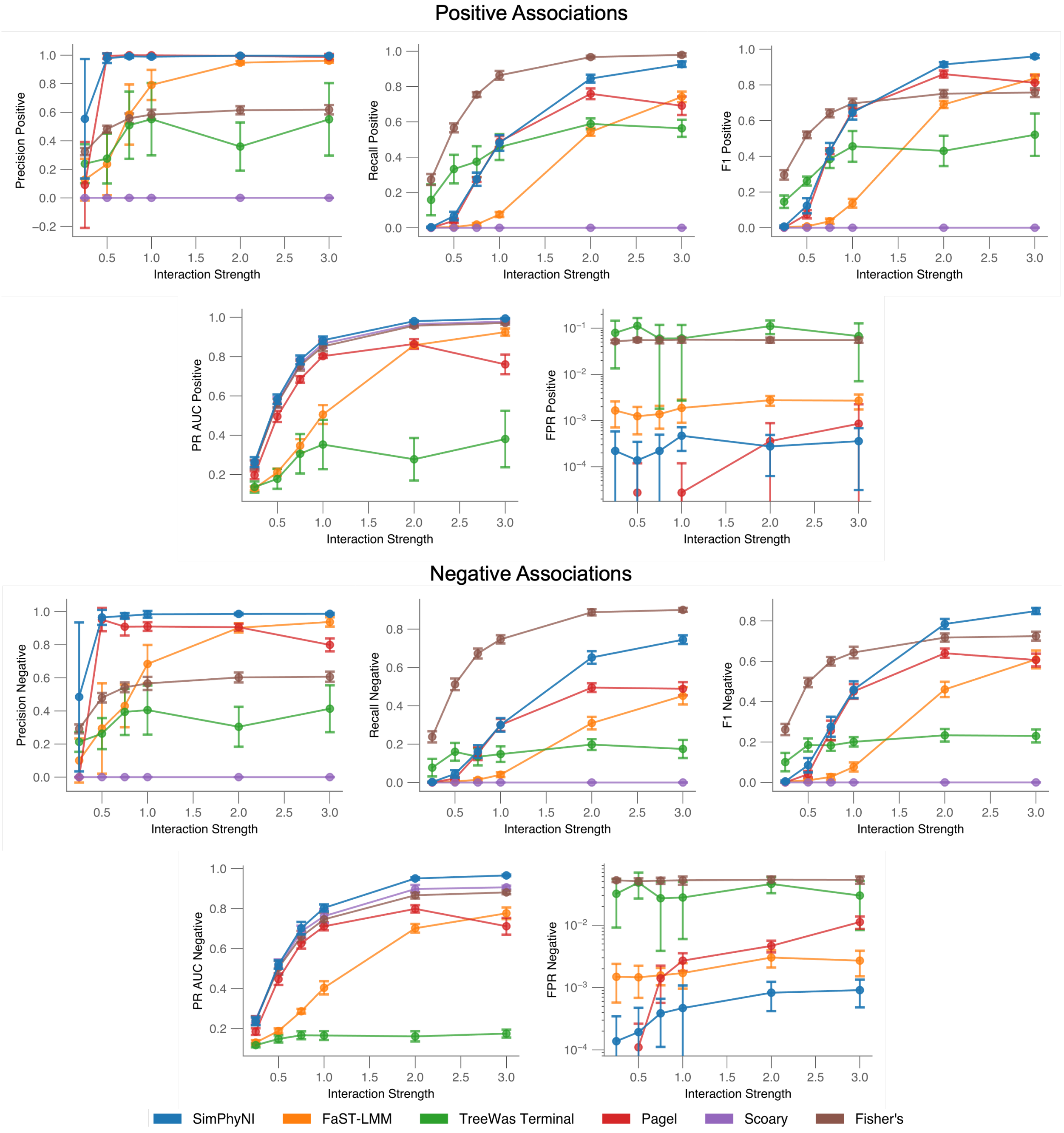


Figure S9: **All metrics for all tested methods using directional 4-state Markov data generation across all tested interaction strengths.** Same format as S8 except data is generated using a directional interaction rather than mutual. A 4-state transition matrix is constructed in which the presence of one trait affects the transition rates of the other, but not vice versa, capturing a model of probabilistic causality. Similar trends to mutual data are seen here with SimPhyNI outperforming other methods.

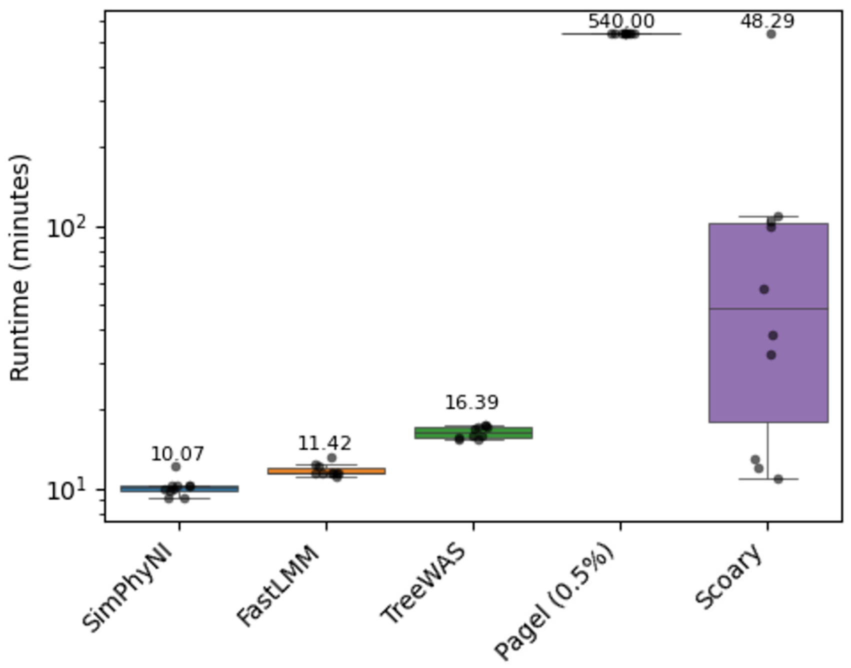


Figure S10: **Runtime of complete workflows for SimPhyNI and other tested methods.** Runtime in minutes for each method on a set of 10 synthetic datasets, each comprised of a synthetic 500 genome phylogenetic tree and 1000 traits. Tools were used to test interactions for all combinations of traits parallelized across 16 CPU cores, with total 32 GB RAM. A time limit of 9 hours was set for analyses with only Pagel's correlation method reaching this limit, completing 0.5% of association tests. Statistical comparison of runtime by Wilcoxon rank sum test shows significantly faster performance of SimPhyNI over all other tested methods S3.

| **Comparison** | **U statistic** | ***p*-value** |
|---|---|---|
| SimPhyNI vs FastLMM | 8.0 | **0.001672** |
| SimPhyNI vs TreeWAS | 0.0 | **0.000181** |
| SimPhyNI vs Pagel | 0.0 | **0.000063** |
| SimPhyNI vs Scoary | 2.0 | **0.000328** |

Table S3: Pairwise statistical comparisons of runtime between **SimPhyNI** and other methods using the Mann–Whitney U test. Bold *p*-values indicate significance ($p < 0.05$).

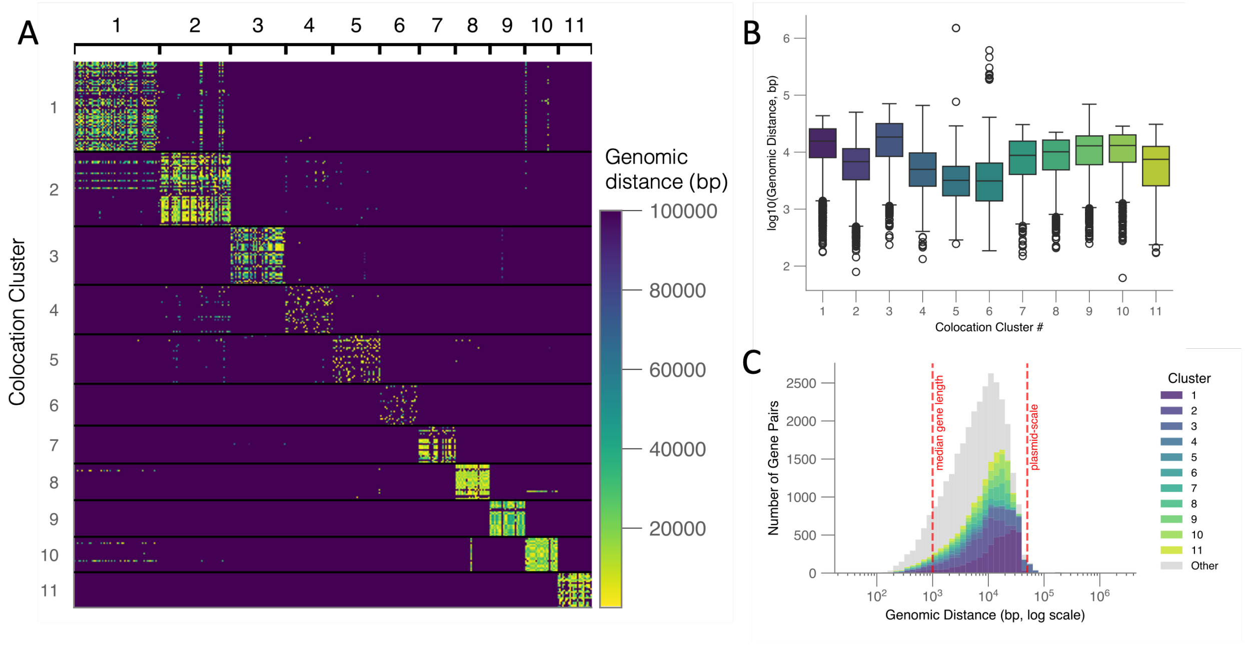


Figure S11: **Genomic distances within co-location clusters made from *E. coli* accessory genes**. (A) A heatmap of genomic distance between genes in the top 11 largest co-location clusters. (B) Distribution of pairwise distances for the top 11 co-location clusters. (C) Within cluster pairwise distances for all co-location clusters. Contributions of top 11 co-location clusters are labeled and reference points for gene and plasmid scale is annotated.

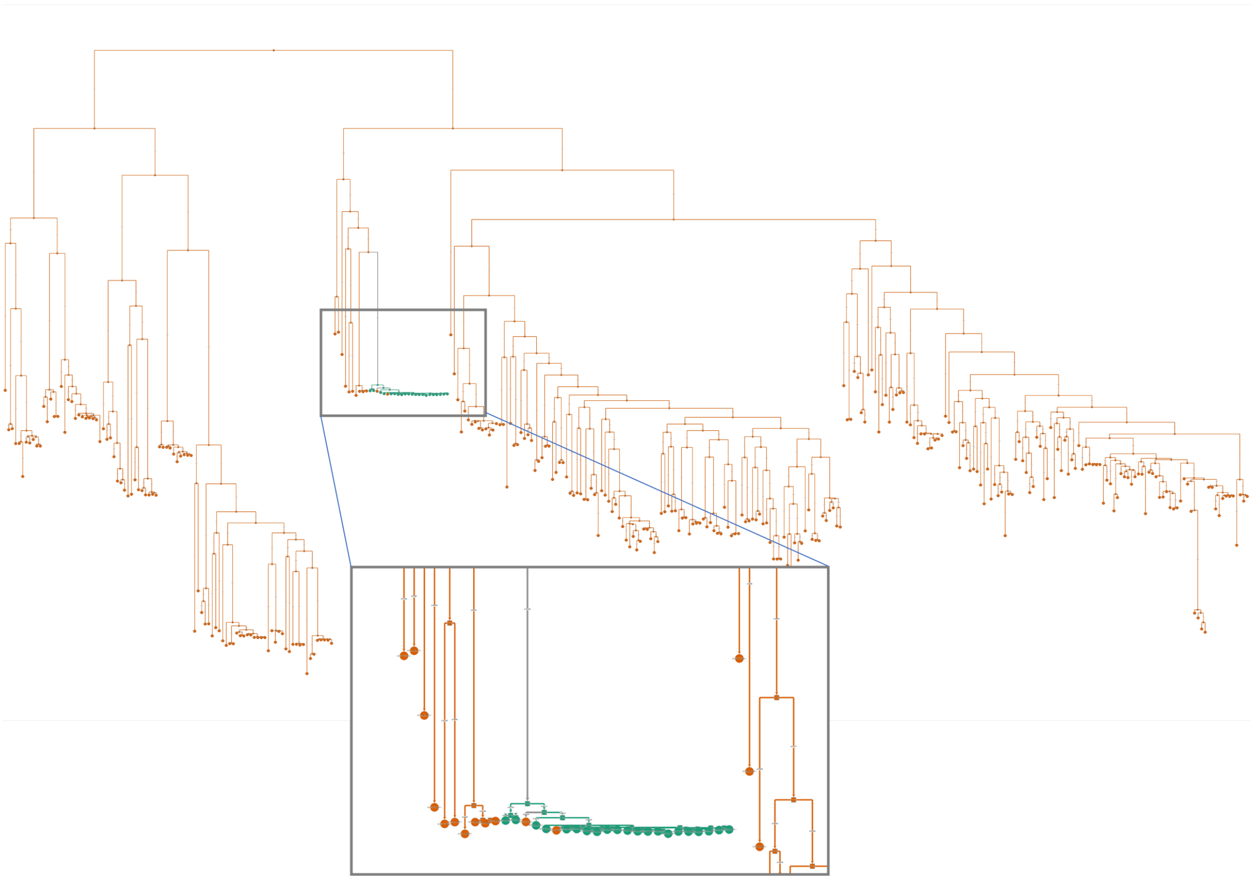

Figure S12: **Poorly resolved ordering in outbreak-like clusters on the phylogenetic tree creates spurious associations.**. A clonal expansion on phylogenetic tree in which multiple events are inferred to occur. Due to near zero genomic distance between strains, the correct ordering of strains becomes ambiguous. Further this outbreak contains the only occurrence of this gene, giving inaccurate transition rates for the gene, introducing spurious associations.

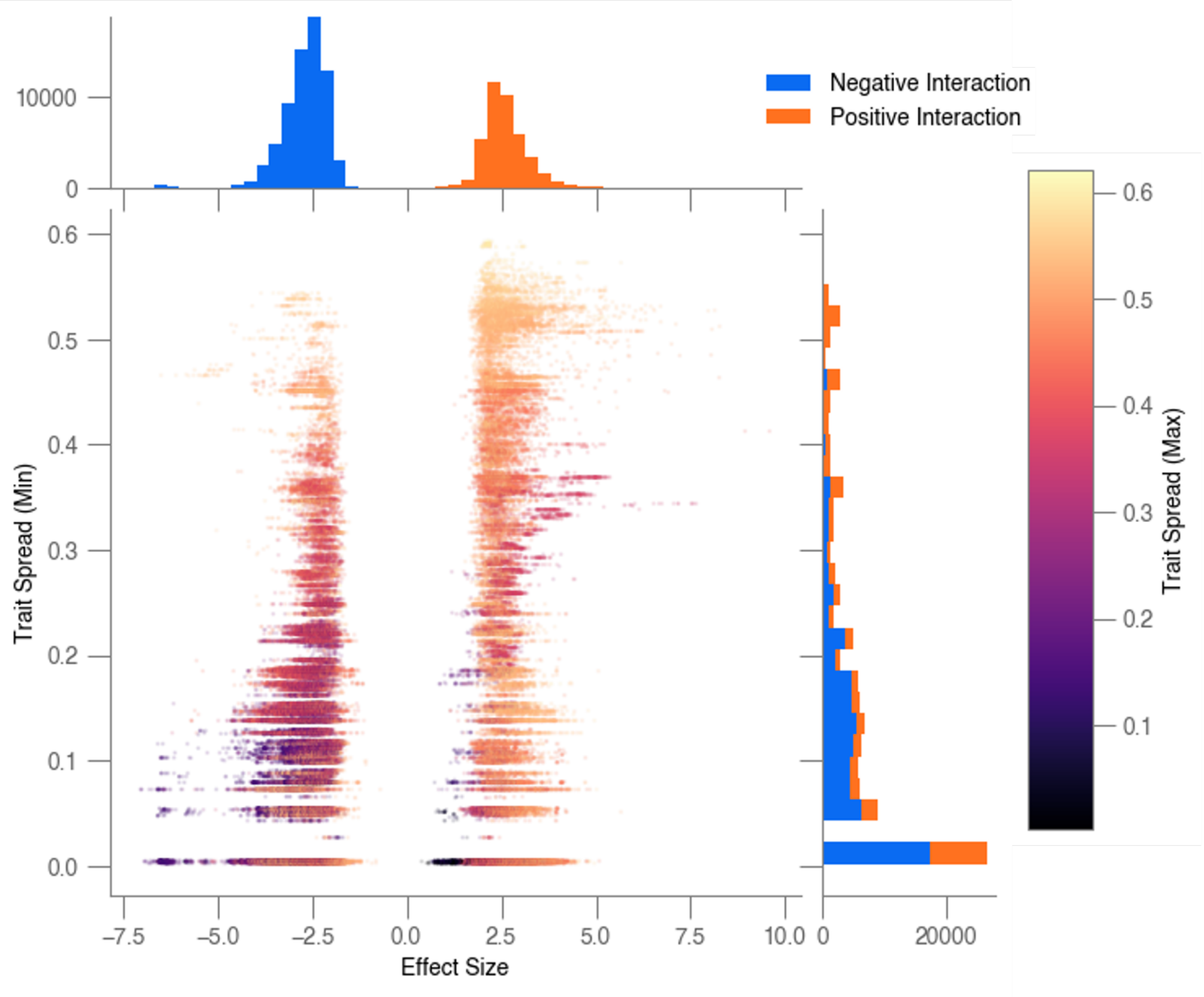


Figure S13: **Distributions of phylogenetic spread compared against effect size of significant interactions between *E. coli* accessory genes**. A scatter plot of significant interactions (Benjamini-Yekutieli $< 0.01$) filtered to remove co-location derived interactions, visualizing their effect size and phylogenetic spread. Position of points is determined by the minimum spread of interacting traits while color is determined by the maximum. Marginals show counts along each axis colored by the direction of the interaction.
